# Diverse signaling proteins recruit RCC1-Like Domain proteins to the plasma membrane through a common structural framework

**DOI:** 10.64898/2026.10.09.757958

**Authors:** Takeshi Nishimura, Yoshinori Hirano, Ana Lopez Vazquez, Martine Trevisan, Peter Sabol, Juan Dong, Toshio Hakoshima, Christian Fankhauser, Miyo Terao Morita

## Abstract

Plants translate diverse environmental cues into spatially regulated growth responses through intracellular signaling mechanisms that remain incompletely understood. Here, we identify the BREVIS RADIX (BRX) domain of RCC1-LIKE DOMAIN (RLD) proteins as a versatile interaction module that recognizes multiple signaling proteins in Arabidopsis. We identified members of the PHYTOCHROME KINASE SUBSTRATE (PKS) and BIG GRAIN (BG) families as novel RLD interactors families, in addition to the previously characterized LAZY1-LIKE (LZY) proteins and BREAKING OF ASYMMETRY IN THE STOMATAL LINEAGE (BASL). Structural, biochemical, and cell-based analyses demonstrate that these diverse partners, despite having divergent interaction sequences, bind a common surface on the RLD BRX domain to recruit RLD to the plasma membrane. This interaction relies on a shared β-strand-based recognition mode, in which distinct peptide conformations accommodate binding to the same BRX-domain surface. Furthermore, we show that *RLD2* and *RLD3* contribute to hypocotyl phototropism, and that disrupting the conserved PKS interaction motif abolishes PKS4 function in this response. Together, our findings identify the RLD BRX domain as a molecular platform for diverse protein interactions and establish a PKS–RLD module that operates in phototropism. Furthermore, these findings strongly suggest that RLD recruitment to the plasma membrane via PKS underlies phototropic signal transduction.

## Introduction

Plants continuously perceive environmental cues such as gravity, light, and water availability and adjust their growth and development accordingly. Various sensing mechanisms, including receptor molecules that detect these environmental signals, have been elucidated. However, the intracellular signaling pathways that link environmental perception to cellular responses remain incompletely understood. Recent studies have identified LAZY1-LIKE (LZY) proteins and RCC1-LIKE DOMAIN (RLD) proteins as key components of gravity-sensing and -signaling in Arabidopsis root gravitropism (1, 2, 3). In root columella cells, LZY proteins localize to amyloplasts and subsequently translocate to the adjacent plasma membrane upon amyloplast sedimentation, thereby establishing polar localization along the gravity vector (2, 4, 5, 6).

RLD proteins, initially characterized as PRAF (PH, RCC1 and FYVE) proteins, have a characteristic multidomain architecture comprising an N-terminal pleckstrin homology (PH) domain, multiple regulator of chromosome condensation 1 (RCC1)-like repeats, an Fab1, YOTB, Vac1 and EEA1 (FYVE) domain and a C-terminal BREVIS RADIX (BRX) domain (7, 8, 9). The PH and FYVE domains of RLD1/PRAF1 bind phospholipids, whereas its RCC1-containing region exhibits guanine nucleotide exchange activity toward Rab GTPases in vitro (7).

The C-terminal conserved CCL motif of LZY physically interacts with the BREVIS RADIX (BRX) domain of RLD proteins, resulting in the recruitment of RLD proteins to the plasma membrane (3). Structural analysis revealed that the LZY CCL motif adopts a β-hairpin structure and binds to a hydrophobic groove in the RLD BRX domain, forming an intermolecular antiparallel β-sheet that extends the three-stranded β-sheet of the BRX domain into a five-stranded β-sheet (3). Functionally, RLD1 and RLD4 regulate lateral root growth direction; the *rld1 rld4* double mutant exhibits an altered expression pattern of the auxin-responsive reporter *DR5rev::GFP* (3). Furthermore, RLD proteins physically interact with the ARF-GEF GNOM, a key regulator of membrane trafficking and auxin transport (10). Polarized RLD proteins are thus thought to regulate auxin transport through GNOM-dependent membrane trafficking, making the interaction between the LZY CCL motif and the RLD BRX domain as a central step in gravity signal transduction (1, 3).

The BRX domain was originally identified as a conserved region of the Arabidopsis BRX protein family and is now recognized as a plant-specific protein–protein interaction module (11). BRX-domain-containing proteins have been implicated in several developmental and signaling processes through interactions with distinct protein partners (3, 10, 12, 13, 14). Genetic studies further suggest broader physiological roles for RLD proteins (3). While the *rld1 rld4* double mutant exhibits defects in gravitropic responses, higher-order mutants display severe developmental abnormalities, indicating that RLD proteins participate in biological processes beyond gravity signaling. Indeed, Arabidopsis RLD proteins interact not only with LZY proteins but also with BREAKING OF ASYMMETRY IN THE STOMATAL LINEAGE (BASL), a polarity factor involved in asymmetric cell division during stomatal lineage development (10, 15). These findings raise the intriguing possibility that the BRX domain mediates interactions with additional, as yet unidentified proteins.

In this study, we screened for additional RLD-interacting proteins and identified diverse BRX-domain-binding factors, including members of the PHYTOCHROME KINASE SUBSTRATE (PKS) family, which function in phototropism (16, 17, 18), and the BIG GRAIN (BG) family, implicated in auxin transport (19, 20). Structural analyses revealed that, despite negligible sequence similarity to the LZY-CCL motif, the BRX-binding motifs of PKS, BG, and BASL share a conserved binding architecture defined by antiparallel β-sheet interactions with the BRX domain. These proteins recruit RLD to the plasma membrane through BRX-domain-mediated interactions, revealing a shared mechanism for RLD recruitment by distinct signaling proteins. Genetic and functional analyses further demonstrated that the interaction between RLD and PKS proteins contributes to phototropic responses. Together, our findings reveal how a common structural interaction module links diverse upstream signaling proteins to RLD recruitment and distinct physiological responses.

## Results

### Identification of proteins interacting with the RLD-BRX domain

Previous studies have shown that RLD proteins participate in gravity signaling and stomatal-lineage development through physical interactions with LZY and BASL, respectively (3, 10). These findings raised the possibility that RLD proteins interact with additional partners to function across a broader range of signaling pathways. To identify proteins that act together with RLDs, we performed a yeast two-hybrid screen using a region of RLD1 spanning the FYVE domain to the C-terminal BRX domain (RLD1-FYVE-BRX) as bait. In addition to LZY family members, the screen recovered a diverse set of candidate interactors. Notably, members of the PHYTOCHROME KINASE SUBSTRATE (PKS) and BIG GRAIN (BG) families were identified as prominent binding partners of RLD1-FYVE-BRX (Extended Table 1 and Extended Data Fig. 1a, b). PKS family proteins act downstream of phototropins to mediate light signaling and are required for normal phototropic response in hypocotyls (16, 17, 18, 21, 22). The BG family protein BG1 has been implicated in the regulation of auxin transport in rice. (23).

We examined the physical association between representative PKS/BG proteins and the FYVE–BRX regions of RLD1 and RLD2, which belong to distinct RLD phylogenic subgroups (3). Both PKS and BG proteins interacted with RLD1 and RLD2, suggesting that this interaction is a general feature of RLD proteins rather than specific to RLD1 (Extended Data Fig. 1b).

Sequence comparison among PKS and BG family members revealed a highly conserved region of approximately 10 amino acids shared between the two families (Extended Data Fig. 1a). This motif was previously designated as Motif D (23).

The consensus sequence of Motif D is SXXS(S/T)DLFE(L/I)E, in which the ninth residue is conserved as isoleucine in the PKS family and leucine in the BG family (23). Importantly, an independent yeast two hybrid screen using a part of PKS4 centred around motif D as bait identified RLD2 and RLD4 as interactors. A series of analyses using yeast two-hybrid system with deleted fragments of AtBG1 and RLD1-FYVE-BRX demonstrate that the Motif D sequence and the BRX domain are necessary and sufficient for the interaction (Extended Data Fig. 1c, d).

### Structural basis for diverse BRX-domain interactions revealed by X-ray crystallography

We previously determined the crystal structure of the RLD2 BRX domain in complex with the LZY3 CCL motif, revealing the structural basis of an interaction central to gravity signal transduction (3). To determine how the BRX domain recognizes other RLD-interacting proteins, we investigated its interactions with AthBG1, PKS1 and BASL using biochemical and structural analyses.

Although Motif D mediates the interaction of AthBG1 and PKS1 with the RLD BRX domain, its sequence shows little similarity to the LZY3 CCL motif. Isothermal titration calorimetry (ITC) showed that Motif D peptides derived from AthBG1 and PKS1 bound the RLD2 BRX domain with dissociation constants (*K_D_*) of 820 nM and 1.23 μM, respectively (Extended Data Fig. 2a, b). BASL, a previously characterized RLD interactor, also bound the RLD2 BRX domain through the previously identified region, with a *K_D_* of 1.83 μM (Extended Data Fig. 2c). We next determined crystal structures of the RLD2 BRX domain in complex with AthBG1 Motif D, PKS1 Motif D and BASL peptides at 2.0, 1.8 and 1.52 Å resolution, respectively (Fig. 1 and Extended Data Table 2). In all three complexes, the BRX domain adopts an essentially same structure consisting of an N-terminal three-stranded antiparallel β-sheet and two C-terminal α-helices, and the bound peptides occupy the same hydrophobic groove between the β3-strand and α1-helix (Fig. 1a–c). The AthBG1 peptide forms an antiparallel β-strand with theβ3-strand of the BRX domain, forming an intermolecular β-sheet, in which Leu321, Phe322 and Leu324 form hydrophobic contacts and Glu323 forms salt bridges with RLD2 Arg1033 (Fig. 1a, d). The PKS1 peptide binds to the same hydrophobic groove of the BRX domain, where hydrophobic interactions and salt bridges contribute to the binding in a manner similar to that observed for AtBG1. Strikingly, PKS1 adopts a distinct conformation, comprising a short β-strand followed by a one-turn α-helix (Fig. 1b, e). The β-strand forms an antiparallel intermolecular β-sheet with BRX, whereas the additional helix makes hydrophobic contacts with the RLD2 α1-helix. Additionally, the main chain of PKS1 Ile287 forms a water-mediated hydrogen bond with that of RLD2 Val1034 (Fig. 1b, e). This helical structure is unique among the BRX-binding peptides examined here.

**Fig. 1.**
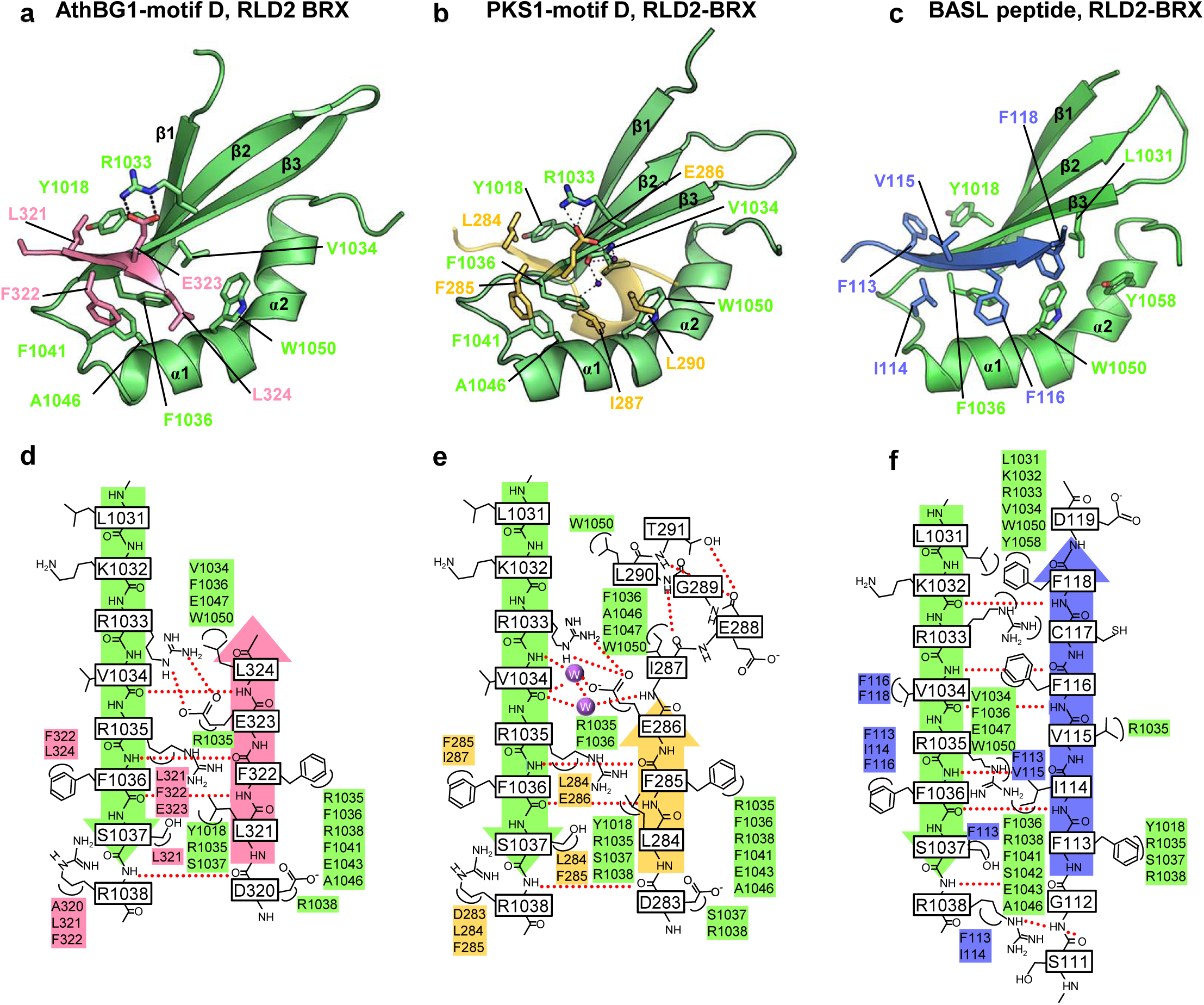
Structural basis of RLD–BRX domain interactions. (a–c) Ribbon representations of crystal structures of the RLD2-BRX domain in complex with (a) AthBG1-motif D (pink), (b) PKS1-motif D (yellow), and (c) BASL peptide (blue). The RLD2-BRX domain (green) comprises a three-stranded antiparallel β-sheet and two α-helices. Dashed lines indicate intermolecular hydrogen bonds. **(d–f)** Schematic representations of the interactions between the RLD2-BRX domain and (d) AthBG1-motif D, (e) PKS1-motif D, and (f) BASL peptide. Hydrogen bonds are shown as dotted lines; van der Waals contacts are also indicated.

The BASL peptide similarly binds the β3–α1 groove and forms an antiparallel intermolecular β-sheet with the β3-strand of the BRX domain (Fig. 1c, f). Unlike AthBG1 and PKS1, however, the BASL peptide is dominated by hydrophobic residues and lacks direct polar contacts with the BRX domain. Thus, despite substantial sequence and conformational differences, AthBG1, PKS1 and BASL engage the same surface of the RLD BRX domain through a shared β-strand-based recognition mode.

To validate the structurally defined interactions, we performed pull-down assays using the RLD2 BRX domain and wild-type or mutated AthBG1, PKS1 and BASL peptides. All of the mutants reduced interaction, supporting the importance of hydrophobic and polar interactions observed in the crystal structures. (Extended Data Fig. 3a–c). Notably, the residues contacting RLD2 Phe1036 were repeatedly identified as major determinants of binding, suggesting a central role for this conserved hydrophobic residue in the recognition of diverse BRX-binding proteins.

### Comparison of binding modes with the LZY3 CCL–BRX complex

Our previous structural study showed that the LZY3 CCL motif adopts a β-hairpin in which both β-strands contact the RLD2 BRX domain (3). To compare BRX-binding peptides within a common structural framework, we designated the six residues of the C-terminal βB-strand of LZY3 as positions 1–6 (P1-P6) (Fig. 2a). In LZY3, a glutamate at P5 is involved in electrostatic interactions whereas the other positions predominantly contribute to hydrophobic interactions. Structural superimposition revealed that the BRX domains adopt essentially same conformations in all complexes, with most interacting residues closely aligned except for Arg1033, whereas the bound peptides exhibit distinct conformations (Fig. 2b–e).

**Fig. 2.**
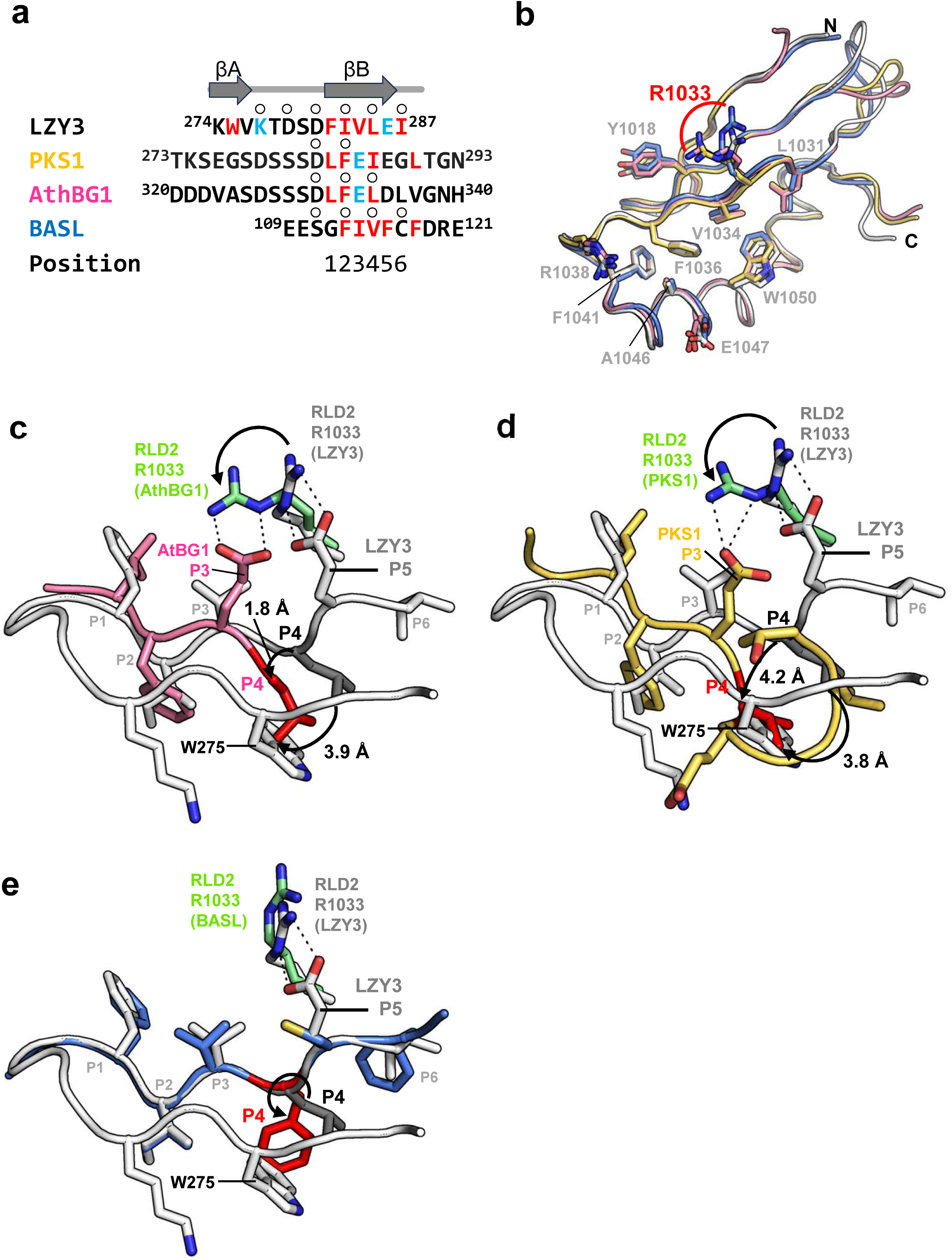
Structural comparison of BRX-domain binding modes. **(a)** Sequence alignment of RLD2-BRX–binding peptides with the secondary structures of LZY3 are shown at the top with β-strands (arrow head). Residues forming the βB-strand are sequentially labeled position 1 to 6 (P1–P6). Residues whose side-chain atoms form intermolecular hydrogen bonds and those making van der Waals contacts are colored cyan and red, respectively; residues whose main chains form hydrogen bonds are indicated by open circles. **(b)** Superimposition of the RLD2-BRX domain bound to AthBG1 (pink), PKS1 (yellow), or BASL (blue) onto that bound to LZY3-CCL (white). Corresponding residues interacting with LZY3-CCL are shown as stick models. **(c)** Overlay of LZY3 (white) and AthBG1 (pink) after superposition of the RLD2-BRX domain bound to AthBG1 (pink) on that bound to LZY3-CCL. Residues forming the βB-strand are sequentially labeled P1–P6. **(d)** As in (c), but showing the RLD2-BRX domain bound to PKS1 (yellow). **(e)** As in (d, e), but showing the RLD2-BRX domain bound to BASL (blue).

In the RLD2–AthBG1 complex, the Motif D peptide forms a shorter β-strand corresponding to the LZY3 βB-strand and retains hydrophobic residues at P1, P2 and P4 (Fig. 2a, c). Glu333 at P3 forms salt bridges with RLD2 Arg1033, whereas LZY3 Glu286 at P5 interacts with RLD2 Arg1033. This alternative interaction is accommodated by reorientation of the Arg1033 side chain. In addition, the P4 residue Leu334 is displaced toward the position occupied by Trp275 in the LZY3 βA-strand, partially compensating for the absence of the βA-strand.

PKS1 employs a related but distinct binding mode (Fig. 2d). Its P3 glutamate interacts with RLD2 Arg1033 in a manner similar to atBG1, whereas the P4 residue Ile287 is displaced further downward than the corresponding residue in AthBG1, resulting in a position that almost completely overlaps with that of LZY3 Trp275 (Fig. 2d). Together with the additional short helical structure of PKS1, this rearrangement provides an alternative hydrophobic interaction pattern with the BRX domain. Additionally, water-mediated hydrogen bonds between the main chains compensate for the limited intermolecular hydrogen bonds provided by the short β-strand (Fig. 1b, e).

The BASL peptide closely resembles the backbone conformation of the LZY3 βB-strand but differs in side-chain organization (Fig. 2a, e). Phe116 at P4 adopts a distinct orientation and partially substitutes for the hydrophobic interactions mediated by LZY3 Trp275. In addition, the P5 glutamate of LZY3 is replaced by Cys117 in BASL, accompanied by repositioning of RLD2 Arg1033 away from the peptide to avoid the unfavorable contacts.

Together, these analyses reveal a common recognition principle for structurally diverse BRX-binding peptides. Although AthBG1, PKS1 and BASL lack the βA-strand present in LZY3 CCL motif, distinct structural rearrangements around P4 compensate for at least part of the hydrophobic interactions normally mediated by LZY3 Trp275 in the βA-strand.

### AthBG1, PKS1, and BASL recruit RLD to the plasma membrane through BRX domain interactions

Previous studies demonstrated that plasma membrane–localized LZY3 recruits RLD proteins via an interaction between the LZY CCL motif and the RLD BRX domain (3). Moreover, BASL, PKS and BG all localize to the plasma membrane (10, 19, 23), a localization that was shown to be important for PKS function (23). We therefore asked whether AthBG1, PKS1, and BASL similarly recruit RLD to the plasma membrane. To test this possibility, we examined the subcellular localization of these proteins and their effects on RLD2 BRX domain (RLD2-BRX) localization in transiently transformed Arabidopsis root protoplasts (Fig. 3 and Extended Data Figs. 4 and 5).

**Fig. 3.**
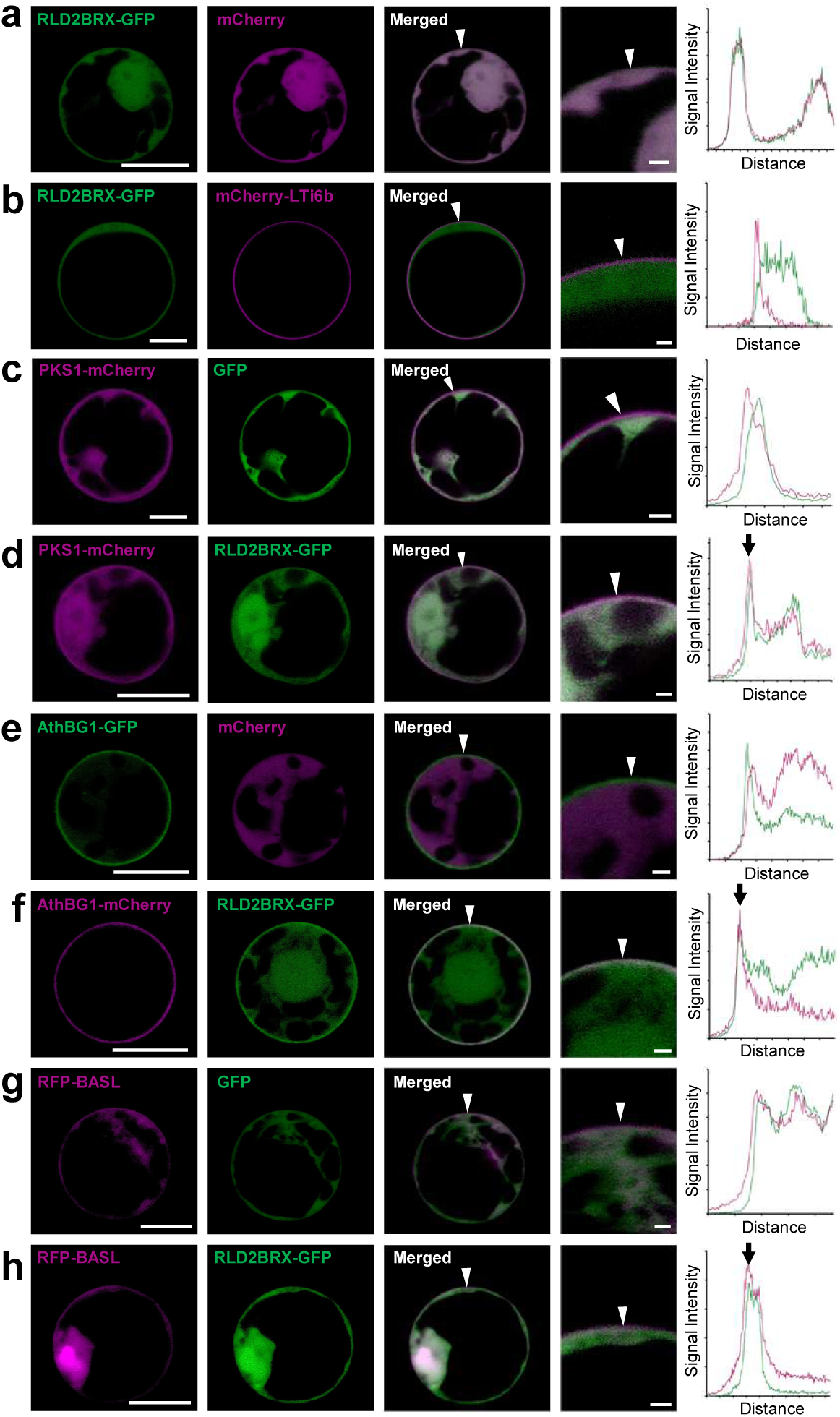
RLD–BRX domain interactions mediate plasma-membrane recruitment and evolutionary diversification. Co-localization analysis of PKS1, AthBG1, and BASL with the RLD2-BRX domain in transiently transformed *Arabidopsis* root protoplasts. Co-expression combinations: (a) RLD2-BRX domain–GFP with cytosolic marker mCherry; (b) RLD2-BRX domain–GFP with plasma-membrane marker mCherry-LTi6b; (c) GFP with PKS1-mCherry; (d) RLD2-BRX domain–GFP with PKS1-mCherry; (e) AthBG1-GFP with cytosolic mCherry; (f) AthBG1-mCherry with RLD2-BRX domain–GFP; (g) BASL-mCherry with GFP; (h) BASL-mCherry with RLD2-BRX domain–GFP. Arrowheads indicate the regions used for intensity plots; arrows indicate overlapping signal peaks at the plasma membrane. Intensity profiles across the ROI are shown in Fig. S4. Scale bar = 10 µm

When expressed alone, RLD2-BRX–GFP localized predominantly to the cytosol, as confirmed using cytosolic mCherry and the plasma membrane marker LTi6b (Fig. 3a, b). In contrast, AthBG1-GFP, PKS1-mCherry, and mRFP-BASL were detected in both the cytosol and at the cell periphery, indicating dual cytosolic and plasma membrane localization (Fig. 3c, e, g, Extended Data Fig. 4). In addition, mRFP-BASL exhibited nuclear localization (Fig. 3g). Co-expression with AthBG1, PKS1, and BASL induced the translocation of RLD2-BRX– GFP to the plasma membrane (Fig. 3d, f, h). Line-scan analysis revealed overlapping fluorescence intensity peaks along the plasma membrane, confirming the co-localization of RLD2-BRX with each interacting partner (Fig. 3d, f, h, Extended Data Fig. 4). Crucially, mutations disrupting the interaction interfaces identified by our structural analysis abolished this membrane recruitment (Extended Data Figs. 3, 4 and 5). mPKS1 carrying F285A/L290A substitutions and mAthBG1 carrying F322A/L324A substitutions, failed to recruit RLD2-BRX– GFP to the plasma membrane (Extended Data Fig. 5a, b). Similarly, BASL-mediated recruitment was markedly reduced by the I114A substitution (Extended Data Fig. 5c). Together, these results provide independent biochemical and cell biological support for the structural models and demonstrate that AthBG1, PKS1, and BASL recruit RLD to the plasma membrane through BRX-domain-mediated interactions. Collectively, these cell-based assays validate our structural models and demonstrate that AthBG1, PKS1, and BASL recruit RLD to the plasma membrane through BRX domain-mediated interactions, analogous to the known LZY-dependent mechanism. Together, these findings indicate that the RLD BRX domain serves as a flexible docking site capable of recognizing multiple protein classes despite low sequence similarity, suggesting that RLDs participate in a broader range of biological processes than previously appreciated.

### PKS proteins require Motif D–mediated interaction with the RLD BRX domain for phototropism

Among the newly identified RLD-interacting partners, the PKS family is of particular interest because PKS members are established regulators of phototropic signaling downstream of phototropins (18, 24). We therefore investigated whether RLD proteins, through their interaction with PKSs, contribute to phototropism. To test this, we examined the phototropic responses of etiolated hypocotyls in various *rld* double-mutant combinations. Among the mutants tested, the *rld2 rld3* double mutant exhibited a significant reduction in phototropic curvature, to an extent comparable to the *pks4-2* mutant (Fig. 4a and Extended Data Fig. 6a–c). In contrast, neither the *rld2* nor *rld3* single mutant displayed a detectable phototropic defect, indicating that *RLD2* and *RLD3* function redundantly in this process. Furthermore, expression of RLD3-mClover3, which complements the developmental defects of the *rld2 rld3* mutant, rescued the phototropic defect, confirming that the phenotype results specifically from the loss of RLD2 and RLD3 function (Extended Data Fig. 6d, e). Because structural and biochemical analyses indicated that PKS proteins bind the RLD BRX domain via Motif D, we next tested whether this interaction is required for phototropism *in vivo*. Point mutations were introduced into Motif D of PKS4 (mPKS4-D1) to generate mutant variants that abolished interaction with RLD proteins while retaining normal subcellular localization (Fig. 4b, c). When expressed in the *pks4-2* mutant background, wild-type PKS4 complemented the phenotype, whereas these Motif D variants failed to restore phototropic responses, demonstrating that Motif D–mediated interaction is required for PKS4 function in phototropism (Fig. 4b, c and Extended Data Fig. 7).

**Fig. 4.**
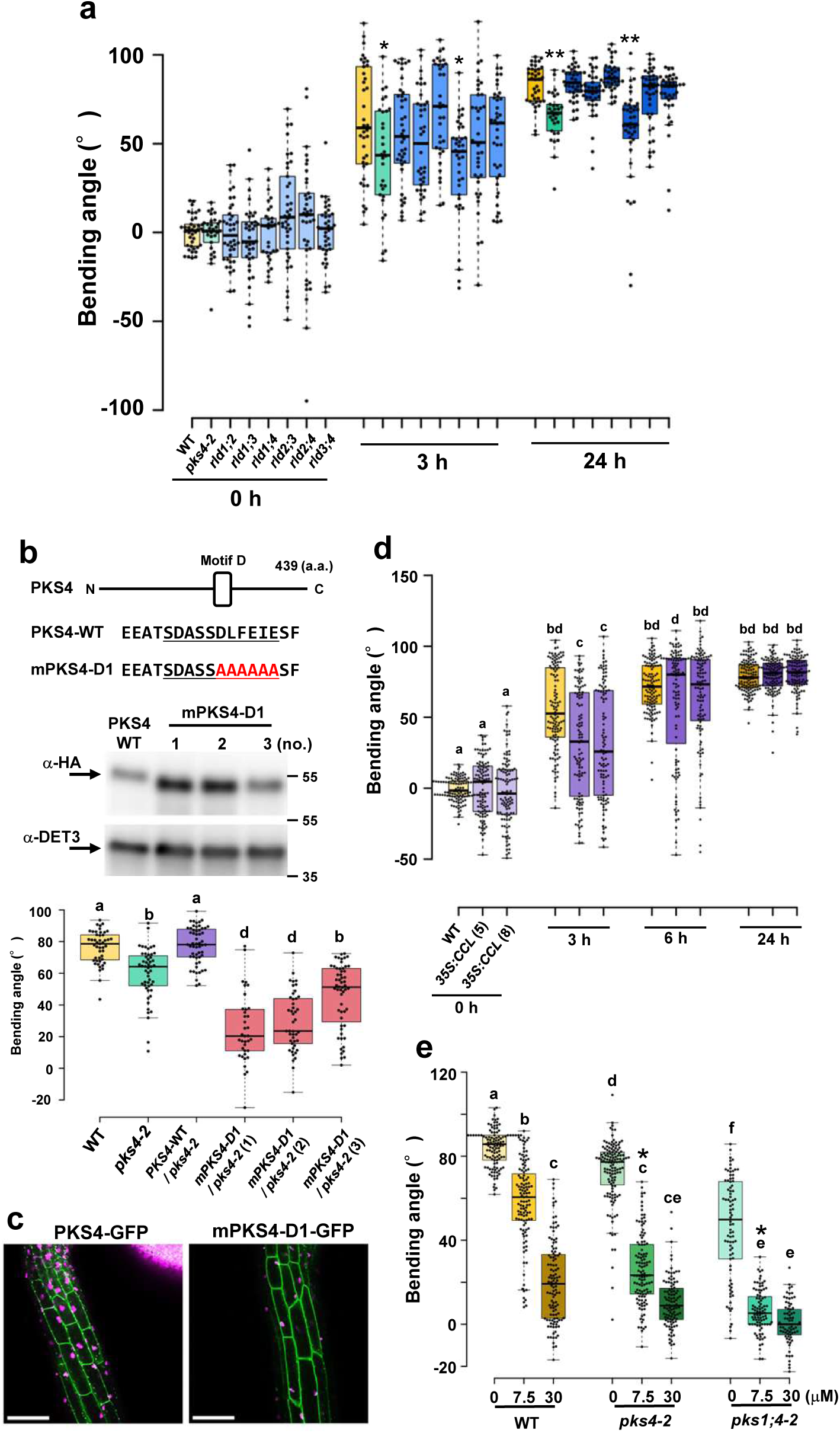
Functional analysis of RLD–PKS interaction in phototropism. **(a)** Phototropism of 4-day-old etiolated hypocotyls of *rld* double mutants with different combinations of *rld1*, *rld2*, *rld3*, and *rld4*. Seedlings were irradiated with unilateral blue light (0.1 μmol m⁻² s⁻¹), and hypocotyl bending was measured after 3 h and 24 h. Curvatures of 30–40 seedlings were quantified. Asterisks indicate significant differences from WT (*, 3 h; **, 24 h) as determined by one-way ANOVA followed by Dunnett’s test. **(b)** Effect of amino-acid substitution in the motif D motif of PKS4 on phototropism. Residues D276–E281 of PKS4 were replaced by alanines (mPKS4-D1), and a C-terminal HA tag was added. The construct was expressed in the *pks4-2* mutant. Phototropic responses were analyzed in lines with confirmed protein expression (Western blot; DET3 as internal control). Etiolated seedlings were irradiated with unilateral blue light (0.1 μmol m⁻² s⁻¹), and curvature was measured after 24 h (n = 37–56). Different letters indicate significant differences (Tukey–Kramer test, *P* < 0.01). Numbers in parentheses indicate independent transgenic lines. **(c)** Subcellular localization of PKS4–GFP and mPKS4-D1–GFP in etiolated hypocotyl cells. Scale bar = 100 µm **(d)** Inhibition of phototropism by overexpression of LZY3-CCL. Seedlings of *35S::CCL-mCherry* were irradiated with unilateral blue light (0.1 μmol m⁻² s⁻¹), and hypocotyl bending was measured after 3 h, 6h and 24 h. Curvatures of 105–119 seedlings were quantified. Different letters indicate significant differences (Tukey–Kramer test, *P* < 0.01). Numbers in parentheses indicate independent transgenic lines. **(e)** Effect of brefeldin A (BFA) on phototropism in *pks4* and *pks1 pks4* double mutants. Four-day-old etiolated seedlings were transferred to media containing the indicated BFA concentrations and continuously irradiated with unilateral blue light (0.1 μmol m⁻² s⁻¹). Curvature was measured after 24 h (n = 78–132). Different letters indicate significant differences (Tukey–Kramer test, *P* < 0.01). A two-way ANOVA revealed a significant genotype × BFA interaction (P = 1.7 × 10⁻²⁴). Planned interaction contrasts further showed that the response to 7.5 µM BFA was significantly greater in *pks4-2* and *pks1 pks4-2* than in WT (Holm-adjusted P = 5.8 × 10⁻¹² and 1.4 × 10⁻⁴, respectively). *Asterisks indicate significant genotype × BFA interaction contrasts comparing the response to 7.5 µM BFA in each mutant with that in WT.

Interestingly, expression of these interaction-deficient PKS4 variants led to a dose-dependent inhibition of phototropism in *pks4-2* plants (Fig. 4b). Similar inhibitory effects were observed with additional point mutations in Motif D (Extended Data Fig. 7b-d). These results indicate a dominant-negative-like effect, further underscoring the critical requirement of an intact Motif D for PKS4 function in phototropic signaling. The LZY CCL motif is the first identified BRX-binding peptide and displays high binding affinity for the RLD BRX domain. To examine whether exogenous expression of a competing BRX-binding motif could perturb phototropism, we analyzed transgenic plants overexpressing the LZY3 CCL motif. Overexpression of LZY3 CCL significantly reduced phototropic curvature 3 hours after unilateral blue-light irradiation, whereas this difference diminished at later time points, indicating a delay rather than a permanent loss of the response (Fig. 4d). These observations are consistent with a model in which excess CCL motif competitively interferes with the PKS–RLD association by occupying the BRX domain.

RLD proteins have been proposed to function together with the sec7 domain protein ARF-GEF GNOM to regulate membrane trafficking (10). To investigate whether sec 7 domain proteins, including GNOM, contribute to PKS-mediated phototropism, we examined the effect of brefeldin A (BFA), a well-characterized inhibitor of GNOM-mediated membrane trafficking pathways (25). BFA treatment inhibited phototropic bending in wild-type seedlings and caused enhanced sensitivity in *pks* mutants (Fig. 4e). Notably, the *pks4-2* and *pks1 pks4* double mutant exhibited stronger BFA sensitivity than the wild type. Together, these findings suggest that GNOM-mediated membrane trafficking is involved in PKS-mediated phototropic signaling. Collectively, our data indicate that the interaction between motif D of PKS proteins and the BRX domain of RLDs constitutes an important step in phototropism and may link phototropic signaling to GNOM-mediated membrane trafficking.

## Discussion

### Structural versatility of BRX domain-mediated interactions

In this study, we demonstrate that the RLD2 BRX domain recognizes diverse peptide sequences derived from LZY, AthBG1, PKS1, and BASL (Figs. 1 and 2). Despite their limited sequence similarity, these proteins employ remarkably similar binding strategies. Structural analyses revealed that all interactors engage the BRX domain through an intermolecular β-sheet formed with the β3 strand of the BRX domain, establishing β-strand-mediated recognition as a central and common feature of BRX domain interactions.

Although AthBG1, PKS1, and BASL bind the BRX domain with comparable micromolar affinities (*K_D_* = 0.82– 1.83 μM), these interactions are substantially weaker than that of the LZY3 CCL motif (*K_D_* ≈ 10 nM) (3). Nevertheless, structural comparisons revealed a surprising mode of convergence (Fig. 2). While the LZY CCL motif adopts a β-hairpin structure in which the N-terminal βA strand contributes a key hydrophobic residue (Trp275), AthBG1, PKS1, and BASL all lack this βA strand. Instead, each of these interactors compensates for the absence of the βA strand through distinct conformational rearrangement at position P4, partially replacing the hydrophobic interactions mediated by LZY3 Trp275. These observations uncover a remarkable structural plasticity in BRX domain recognition and explain how the BRX domain accommodates multiple structurally and functionally distinct partners within a shared interaction framework.

Interestingly, AthBG1 displays a higher binding affinity for the BRX domain than PKS1, despite only five residues being well-ordered in the crystal structure of RLD2 BRX domain complexed with AtBG1. Furthermore, truncating regions outside this resolved core markedly reduced binding affinity. These findings suggest that structurally flexible or disordered regions flanking the core β-strand contribute to complex formation, likely through transient interactions or entropic effects. Alternatively, crystal packing effects may have influenced the observed conformation of the AthBG1 peptide, particularly at its C-terminal region. Future structural analyses will be required to distinguish between these possibilities.

### Recruitment of RLDs by diverse plasma membrane-associated proteins

A key common feature shared by all currently identified BRX-binding factors is their localization to the plasma membrane (2, 5, 6, 10, 19, 23). Previous studies established that LZY proteins recruit RLDs to the plasma membrane via an interaction between the LZY CCL motif and the RLD BRX domain (3). Here, we show that AthBG1, PKS1, and BASL similarly recruit the RLD BRX domain to the plasma membrane in a motif-dependent manner (Fig. 3).

These findings suggest that the targeted recruitment of RLD to specific plasma membrane domains represents a general consequence of BRX-mediated interactions. Although plasma membrane enrichment of AthBG1, PKS1, and BASL was detectable in our protoplast assays, their membrane signal intensity appeared less pronounced than that of LZY proteins (Fig. 3). One possibility is that these proteins differ in their intrinsic lipid-binding preferences or membrane-association modes, which may require specific cellular contexts or lipid compositions not fully preserved in isolated protoplasts (23).

Notably, several BRX-binding proteins exhibit distinct polarized localization in planta. For instance, LZY proteins accumulate on the lower (gravity-facing) plasma membrane in columella cells, whereas BASL establishes cortical polarity during asymmetric cell division (2, 10). Such localized distribution is thought to spatially restrict downstream signaling events. In contrast, polarized localization has not been reported for PKS proteins (18, 23). Because directional light cues must ultimately be translated into asymmetric responses during phototropism, an important open question is whether PKS proteins, RLD proteins, or upstream phototropin-signaling components undergo light-dependent subcellular redistribution or polar localization. Based on these observations, we propose a unifying model in which functionally distinct, plasma membrane-associated signaling proteins recruit RLDs via their respective BRX-binding motifs, thereby positioning RLDs at defined membrane domains in response to specific developmental or environmental cues (Fig. 5a). This mechanism would allow diverse upstream signaling pathways to access RLD-dependent cellular outputs while preserving spatial specificity.

**Fig. 5.**
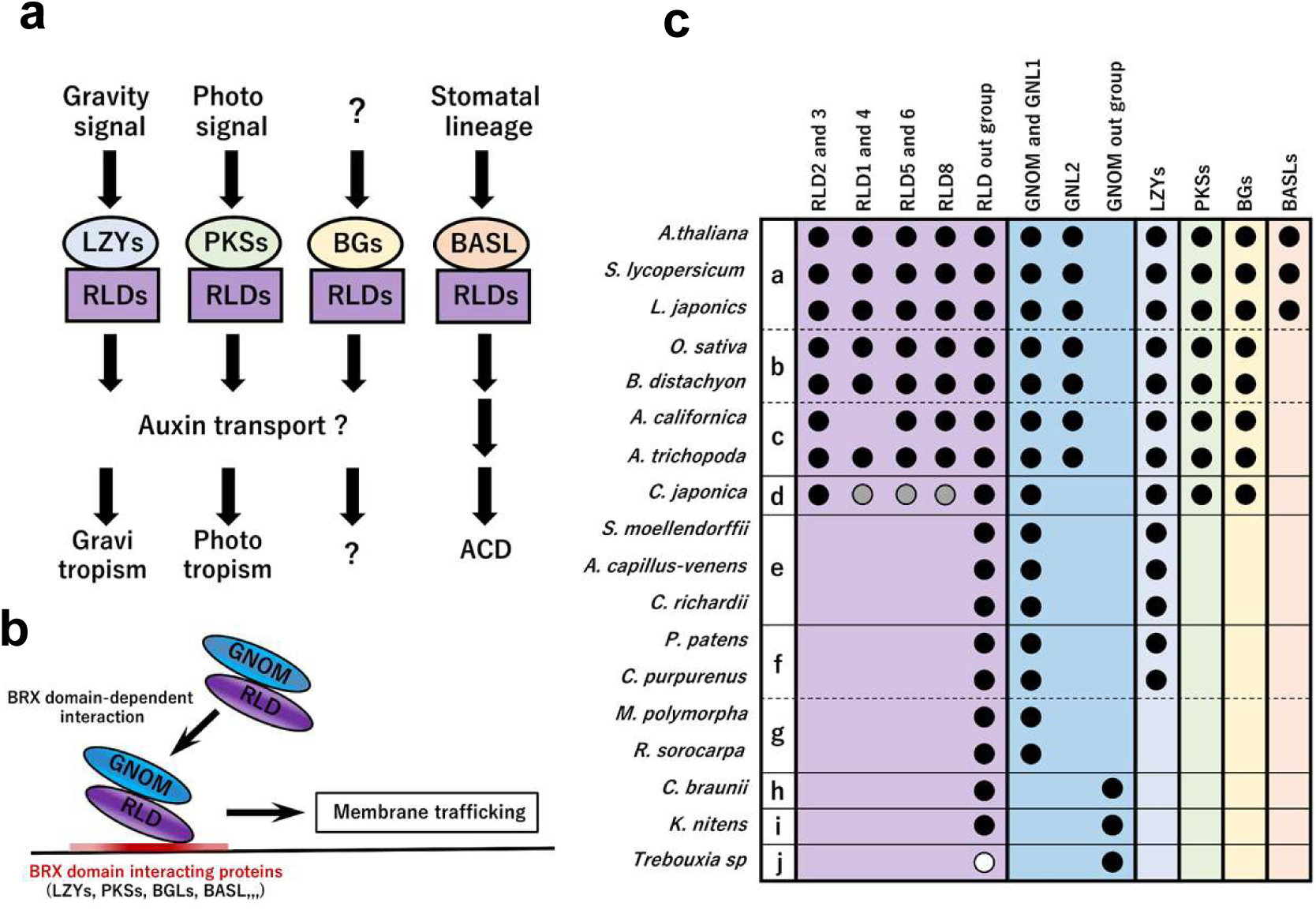
Model and evolutionary diversification of RLD-associated signaling. **(a)** Model illustrating interactions of RLD with multiple partners, including LZY, PKS, AthBG, and BASL, through the BRX domain and their involvement in distinct physiological processes. **(b)** Model illustrating the intracellular dynamics of RLD. LZY, PKS, AthBG, and BASL localize to the plasma membrane and recruit RLD through BRX-domain-mediated interactions. RLD also interacts with GNOM, which is involved in the regulation of membrane trafficking. **(c)** Presence and diversification of RLD, GNOM, LZY, PKS, AthBG, and BASL families across green algae and land plants (Viridiplantae). Homologs were identified by BLAST searches against the NCBI database. All sampled species belong to Streptophyta except *Trebouxia* sp. (j, Chlorophyta). Groups are classified as follows: (a) eudicots; (b) monocots; (c) magnoliids + Amborellales; (d) gymnosperms; (e) ferns + lycopods; (f) mosses; (g) liverworts; (h) Charales; (i) Klebsormidiales; and (j) Trebouxiophyceae. RLD and GNOM subfamilies were classified based on the phylogenetic analyses shown in Extended Data Fig. 8. Black circles indicate the presence of homologs, gray circles indicate closely related homologs, and white circles indicate proteins with low overall sequence similarity but conservation of key functional domains.

Among the proteins functionally associated with RLDs, the ARF-GEF GNOM is particularly intriguing because it physically interacts with RLDs and serves as a central regulator of endosomal trafficking and auxin transport (10, 25, 26). Although the molecular actions of GNOM at or near the plasma membrane remain incompletely defined (27), its association with RLDs suggests that the recruitment of RLD by diverse signaling modules provides a mechanism to spatially couple upstream signal to GNOM-associated cellular processes (Fig. 5b). Rather than implying that all BRX-binding proteins regulate an identical downstream output, this model suggests that RLD acts as a shared molecular hub through which spatially restricted cues engage membrane-associated cellular functions.

Previous study has shown that the plasma membrane localization of the AGC kinase D6PK relies on GNOM-mediated trafficking (28, 29). Furthermore, the polarized localization of LZY in gravity-sensing columella cells directs the local enrichment of D6PK (6). Integrated with our findings, these insights raise the compelling possibility that special positioning of RLD-interacting proteins organizes a core membrane-associated signaling complex comprising RLD, GNOM, and D6PK. Such a mechanism could provide the long-sought molecular bridge between environmental signal perception (e.g., gravity and light) and PIN-mediated polar auxin transport, thereby driving directional auxin redistribution during both gravitropism and phototropism.

### The PKS–RLD module contributes to phototropic signaling

A major finding of this study is that RLD proteins function not only in gravitropism but also in phototropism (Fig. 4). We demonstrated that the *rld2 rld3* double mutant exhibits a reduced phototropic bending response comparable to that of *pks4* mutants, establishing a novel role for RLD proteins in phototropic signaling. Furthermore, disruption of the Motif D sequence abolished PKS4 function in phototropism without affecting its subcellular localization, indicating that physical interaction with the RLD BRX domain is strictly required for PKS function in vivo.

The functional importance of the PKS–RLD interaction is further supported by our findings with the LZY3 CCL motif. Because the CCL motif binds the BRX domain with nanomolar affinity, ectopic expression of the CCL peptide is predicted to outcompete endogenous BRX-binding partners for domain occupancy (3). Consistent with this prediction, overexpression of LZY3 CCL motif delayed phototropic bending, supporting the model in which RLD binding to PKS via BRX-domain is critical for PKS-mediated light signaling.

Although the phototropic phenotype of *rld2 rld3* double mutant was relatively modest, similar weak or partial phenotypes have been reported for specific RLD mutant combinations in root gravitropism (3). Because higher-order *rld* mutants display severe overall developmental defects, dissecting higher-level genetic redundancy remains challenging (3). Future studies focusing on the spatiotemporal dynamics of PKS and RLD proteins upon unilateral light stimulation will be essential to elucidate how directional light cues are transduced into asymmetric growth responses.

The dose-dependent inhibition of phototropism induced by Motif D-mutated PKS4 variants are particularly noteworthy. This inhibitory effect likely reflects a dominant-negative mechanism. One plausible model is that these mutant PKS4 proteins retain binding to upstream signaling components (such as phototropins or light-signaling factors) while losing their ability to engage RLDs, thereby sequestering upstream machinery and interfering with endogenous PKS signaling. Further mechanistic studies will be required to test this possibility; nonetheless, these observations reinforce the indispensability of an intact Motif D for PKS function.

The enhanced sensitivity of *pks* mutants to brefeldin A suggests that GNOM may be involved in PKS-mediated signaling during phototropism. Although the mechanistic relationship among PKS, RLD, and GNOM remains to be established, these findings raise the possibility that PKS–RLD interactions are functionally linked to GNOM-dependent membrane trafficking. These results further provide strong support for our proposed model, in which RLDs serve as a molecular bridge linking environmental signal perception and polar auxin transport.

### Evolutionary diversification of RLD-associated signaling pathways

Our phylogenetic analyses provide insight into the evolutionary history of RLD-associated signaling pathways (30). Homologs of GNOM and RLD were identified in green algal lineages as well as in land plants (Viridiplantae), suggesting that the evolutionary origins of these protein families predate the emergence of land plants (Fig. 5c and Extended Data Fig. 8). In contrast, known BRX-binding proteins appeared at distinct stages during land plant evolution. LZY family members are first detected in bryophytes, whereas PKS and BG family proteins emerged later in vascular plants, and BASL appears to be restricted to dicotyledonous lineages.

These observations raise the possibility that ancestral RLD and GNOM proteins originally functioned in fundamental cellular processes prior to the emergence of the diverse BRX-binding adaptors found in extant land plants. During the evolution of land plants, the progressive acquisition of novel BRX-binding proteins may have connected RLD-associated cellular machinery to newly evolved developmental and environmental signaling pathways (Fig. 5a). The accompanying diversification of the RLD family may have further expanded the spatial specificity and regulatory dynamics of these interactions. Thus, RLD proteins may have flexibly incorporated new upstream signaling partners across evolutionary time while preserving the BRX domain as a conserved interaction platform.

It is particularly intriguing that Motif D is highly conserved across both the PKS and BG families. The preservation of this common BRX-binding module suggests that interaction with RLDs may represent an evolutionarily conserved feature of both lineages. Although the physiological roles of BG proteins remain poorly understood, the strong conservation of Motif D suggests that its interaction with RLDs may have functional significance. Determining the physiological role of the BG–RLD interaction will therefore be important for understanding how RLD-associated signaling pathways diversified during plant evolution.

## Concluding remarks

Collectively, our findings reveal an unexpected diversity of proteins that engage the RLD BRX domain. Despite extensive sequence divergence, these partners share a conserved structural framework for BRX recognition and function to recruit RLDs to the plasma membrane. Furthermore, functional characterization of the PKS– RLD module demonstrates that BRX-mediated interactions directly drive phototropic signaling. We therefore propose that RLD proteins serve as versatile signaling platforms that have been repeatedly deployed across distinct physiological pathways through interactions with diverse membrane-associated factors. A general consequence of these interactions may be the targeted recruitment of RLD-associated machinery to defined plasma membrane microdomains. Given the emerging central role of GNOM in RLD-dependent signaling pathways, deciphering how RLDs dynamically direct the spatial regulation of GNOM activity represents a key challenge for future research. Such studies will ultimately illuminate how diverse environmental and developmental inputs converge on shared cellular modules to coordinate plant growth and environmental adaptation.

## Acknowledgments

We are grateful to Masahiko Furutani and Hiromasa Shikata for helpful discussions. We also thank Wakana Takase, Mai Naganuma, Yuka Yamada and Yuri Komada for technical assistance. We appreciate access to BL41XU beamlines at the SPring-8 synchrotron facility and BL-1A beamline at the Photon factory for the provision of synchrotron data-collection facilities (proposal nos.2016A2519, 2016B2519, 2017B2552, 2018A2540, 2018A2503 and 2022A2766). This work was supported by a Core Research for Evolutionary Science and Technology (CREST) award from the Japan Science and Technology Agency (JST) JPMJCR14M5 to M.T.M., a MEXT Grant-in-Aid for Scientific Research on Innovative Areas 18H05488 to M.T.M., a MEXT Grant-in-Aid for Transformative Research Areas JP26H00449 to M.T.M, the Mitsubishi Foundation to M.T.M., a Japan Society for the Promotion of Science Grants-in-Aid for Scientific Research (KAKENHI) Grant Numbers JP23K05826, JP24H00855 and JP26K02055 to T.N. JP25K09523 to Y.H, Grants from the Swiss National Science Foundation (310030B_179558 and 320030-236083) to C.F. We thank the Cellular Imaging Facility (https://cif.unil.ch/) for help with microscopy. This work was also supported by Optics and Imaging Facility, the Model Plant Section of Model Organisms Facility, and Trans-Scale Biology Center in National Institute for Basic Biology.

## Author contributions

T.N. conceived and designed the study, collected and analyzed the data, and wrote the manuscript. Y.H. performed the crystallographic and biochemical analyses and contributed to writing the manuscript. A.L.V., M.T. and P.S. acquired, analyzed and interpreted the phototropism data. J.D. provided unpublished information related to BASL. T.H. contributed to the design of the crystallographic analysis. C.F. conceived and supervised the analysis of the PKS–RLD interaction and its functional role in phototropism, analyzed and interpreted the data, and revised the manuscript. M.T.-M. conceived and designed the study, supervised the research, and contributed to writing the manuscript.

## Data availability

The coordinates and structure factors for the RLD2-BRX–AthBG1-Motif D, RLD2-BRX–PKS1-Motif D and RLD2-BRX–BASL complexes have been deposited in the Protein Data Bank under accession codes 46PV, 46PW and 46PX, respectively. All other data supporting the findings of this study are available from the corresponding author upon reasonable request.

## Materials and Methods

### Plant materials and growth conditions

Arabidopsis thaliana (L.) Heynh. accession Columbia-0 (Col-0) was used as the wild type. The following mutant alleles and transgenic lines were used in this study: *rld1-2*, *rld2-2*, *rld3-2*, *rld4-1* (3), *pks1-1*, *pks4-2* (18), and *PKS4p::PKS4-GFP* (23). The primers used in this study are listed in Supplementary Table 3. Surface-sterilized seeds were sown on Murashige and Skoog (MS) agar medium containing 1× MS salts, 1% (w/v) sucrose, 0.01% (w/v) myo-inositol, 0.05% (w/v) 2-(N-morpholino) ethanesulfonic acid (MES), and 0.5% (w/v) gellan gum, adjusted to pH 5.8. Seeds were stratified in the dark at 4 °C for 3 d and subsequently grown vertically at 23 °C under continuous white light (80–120 μmol m^−2^ s^−1^) for 10–14 d. Seedlings were transferred to soil when required.

To generate *RLD3p::RLD3-mClover3*, the mClover3 coding sequence was amplified from pNCS-mClover3, which was kindly provided by Michael Lin (Addgene plasmid #74236; (31)), and inserted upstream of the nopaline synthase terminator in a pENTR vector using In-Fusion cloning (Takara Bio), generating pENTR-mClover3. A 1,017-bp fragment corresponding to the RLD3 promoter region, spanning −1,132 to −115 bp relative to the translational start site of the RLD3 transcript (NM_121947.4), was amplified from Col-0 genomic DNA and inserted upstream of the mClover3 coding sequence. The RLD3 coding sequence was then inserted between the RLD3 promoter and mClover3, generating pENTR-*RLD3p::RLD3-mClover3*. The resulting entry clones were transferred into the binary vector pGWB501 using LR Clonase II (Thermo Fisher Scientific).

The cloning strategy to generate PKS4 variants used in this study is based on a previous report (23). The coding sequence of PKS4 motif D of construct *PKS4p::PKS4-3XHA* (pPS09) was changed by ordering modified coding sequences of PKS4 (from Eurofins) and replacing them in construct pPS09. We obtained mPKS4-D1 in which the core DLFEIE was mutated into AAAAAA (pAL09), mPKS4-D2 core DLFEIE was mutated into ALFAIE (pAL35) and mPKS4-D3 core DLFEIE was mutated into DLAAIE (pAL36). *PKS4p::PKS4-GFP* was described in previous report (23). To generate the mPKS4-GFP in which the core DLFEIE was mutated into AAAAAA we used the same strategy as described above to obtain construct.

### Generation of *rld2* and *rld3* CRISP alleles

To generate *rld2; rld3* double mutant by CRISPR/Cas9 we selected 3 targets in RLD2 (At5g12350) and RLD3 (At5g19420) using CRISPR-P: target1 AAAAGTAAACAACTGCCCAG (RLD3), target2 TCTTGGGGAGAGGAATCTGA (RLD2 and RLD3), target3 ATAGCCCAAGAACATATACC (RLD2). These 3 targets were cloned into pEC1.2/EC1.1_spCas9_GFP (Addgene plasmid # 161933) using the protocol described in previous report (32) to give pMT46.

We transformed pMT46 construct into *Agrobacterium* strain GV3101/2. We transformed *Arabidopsis* Col-0 wild-type plants with floral dipping. We screened the collected seeds from the T0 plants for GFP-seeds fluorescence and the GFP-positive seeds were sown on ½ × MS plates. We extracted genomic DNA from T1 transgenic plants. We analyzed the presence of mutations by amplifying fragments surrounding the target sites and Sanger sequencing. Interesting T1 plants were transferred on soil and we selected a double homozygous mutant in T2 and named it *rld2-101 rld3-102*. It was then backcrossed to Col-0 and in F2 we selected single homozygous *rld2-101*, single homozygous *rld3-102* and double homozygous *rld2-101; rld3-102*. *rld2-101* has a deletion of 26 pb after the base 996 of the CDS. This leads to an early stop after 335aa of the 1075aa *RLD2* protein. *rld3-102* has an insertion of a G after the base 1410 of the CDS. This leads to an early stop after 476aa of the 1139aa *RLD3* protein.

### Yeast two-hybrid screening and interaction assays

Yeast two-hybrid screening assay for RLD interactors was performed using the Matchmaker Gold Yeast Two-Hybrid System (Clontech/Takara Bio). For library screening, the coding region corresponding to RLD1-FYVE-BRX (CDS positions 1891–3234) was cloned in frame downstream of the GAL4 DNA-binding domain in pGBKT7. The resulting bait plasmid was introduced into the Y2HGold yeast strain using the Fast-Yeast Transformation Kit (G-Biosciences). Library screening was performed using the Mate & Plate Universal Arabidopsis Normalized Library (Clontech/Takara Bio) according to the manufacturer’s instructions.

Positive colonies that grew on synthetic dropout medium lacking Leu and Trp and containing 40 μg/ml X-α-Gal (Glycosynth) and 125 ng/ml aureobasidin A (Takara Bio) were selected and restreaked on synthetic dropout medium lacking Leu, Trp, His and Ade and containing X-α-Gal and aureobasidin A. Blue colonies that grew under these selective conditions were isolated. Plasmids were recovered from yeast colonies and sequenced. Genes identified from multiple independent yeast colonies were selected as candidate RLD1-FYVE-BRX-interacting proteins and are listed in Extended Table 1.

A yeast two-hybrid (Y2H) screening for PKS4 interactors was performed by Hybrigenics Services (Evry, France) using the proprietary ULTImate Y2H™ platform. The sequence comprising amino acids 250 to 308 (centered on motif D) of PKS4 was cloned into the bait vector and screened against an Arabidopsis light-grown seedling cDNA library (146 million interactions were tested). Positive clones were isolated on selective media, sequenced, and identified by comparison with the Arabidopsis genome annotation. Candidate interactors were ranked according to the Predicted Biological Score (PBS) assigned by Hybrigenics. The most promising interactions were tested by follow-up yeast two-hybrid interactions in our laboratory using the WT PKS4 bait and a mutant version of PKS4 in which the conserved sequence of motif D (DLFEIE) was mutated into AAAAAA. Clones that interacted with the WT PKS4 bait but did not interact with mutant PKS4 bait were considered for subsequent analyses.

For directed interaction assays, coding sequences of full-length AthBG family proteins (AthBG1, 1086 bp; AthBG2, 1023 bp; AthBG3, 1098 bp; AthBG4, 1071 bp), full-length PKS family proteins (PKS1, 1317 bp; PKS2, 1326 bp; PKS3, 1161 bp; PKS4, 1218 bp), RLD fragments including RLD1-FYVE-BRX and RLD2-FYVE-BRX (CDS positions 1963–3186), and AthBG1 fragments including AthBG1-Motif D (CDS positions 970–1011) and AthBG1ΔCter (CDS positions 1–1011) were amplified by PCR. DNA fragments were cloned into pGBKT7 or pGADT7 to generate fusion proteins with the GAL4 DNA-binding domain or GAL4 activation domain, respectively.

Bait and prey plasmids were introduced into Y2HGold and Y187 yeast strains, respectively. Auto-activation was tested for all bait constructs before interaction assays. After mating, diploid yeast cells were spotted onto synthetic dropout medium lacking Leu, Trp, His and Ade and incubated at 30 °C for 3 d to assess protein– protein interactions.

For analysis of PKS4 Motif D variants, we used a bait centered on motif D corresponding to amino-acids 250-308 of PKS4. PKS4 wild-type and mutants mPKS4-D1–D6 were generated and cloned into pGBKT7. RLD2 and RLD4 were cloned into pGADT7. Interactions were examined by growth on synthetic dropout medium lacking Leu, Trp and His supplemented with 5mM 3-amino-1,2,4-triazole, and by reporter activation on medium containing β-Gal.

### Phototropism assays

For etiolated growth, seedlings were grown vertically on agar plates in complete darkness at 22 °C for 3 d. For phototropism assays, seedlings were exposed to continuous unilateral blue light (0.1 μmol m^−2^ s^−1^) at 22 °C for up to 24 h.

For brefeldin A (BFA) treatment, 3-d-old etiolated seedlings were transferred to agar medium supplemented with the indicated concentrations of BFA, acclimated for 2 hours in darkness and exposed to continuous unilateral blue light (0.1 μmol m^−2^ s^−1)^. Hypocotyl curvature was quantified 24 h after the onset of irradiation. Images were acquired at the indicated time points using an infrared CCD camera system. Hypocotyl curvature was quantified as the deviation of the mean orientation angle of the upper hypocotyl region, defined as 85– 95% of total hypocotyl length, from the vertical axis after blue-light irradiation. Measurements were performed using a custom MATLAB script developed in the Fankhauser laboratory or the Fiji angle measuring tool. One-way ANOVA (aov) and Compute Tukey’s Honest Significance Differences (HSD.test) [agricolae package] using R software was performed. We considered p-values < 0.01 (0.05 in T1 lines) significant.

### Transient expression assays in Arabidopsis root protoplasts and microscopy

The constructs *35S::mCherry*, *35S::GFP*, *35S::mCherry-LTi6b*, *35S::GFP-LTi6b*, and *35S::RLD2-BRX* domain-GFP were described previously (2, 3). Full-length coding sequences of AthBG1 and PKS1 were amplified by PCR from Col-0 genomic DNA. The full-length BASL coding sequence was amplified from cDNA synthesized from total RNA isolated from Col-0 seedlings. The AthBG1, PKS1 and BASL coding sequences were cloned downstream of the cauliflower mosaic virus 35S promoter in pUC19-based vectors containing either GFP or mCherry or mRFP and the nopaline synthase terminator using In-Fusion cloning (Takara Bio). Point-mutated versions of PKS1, AthBG1 and BASL were generated by PCR using primers containing the desired nucleotide substitutions. The resulting fragments were inserted downstream of the 35S promoter in *pUC19-35S::mCherry* or *pUC19-35S::mRFP* using In-Fusion cloning. The two fragments were assembled and cloned downstream of the 35S promoter in *pUC19-35S::mCherry* or *pUC19-35S::mRFP* using In-Fusion cloning.

For Preparation and transfection of root protoplasts, roots were excised from five 8–9-d-old Arabidopsis Col-0 seedlings and transferred to 2-ml tubes containing 1.5 ml of enzyme solution comprising 1% (w/v) Cellulase, 0.25% (w/v) Pectolyase, 0.4 M mannitol, 10 mM CaCl_2_, 20 mM KCl and 20 mM MES (pH 5.7). Samples were gently rotated at 25 °C for 1 h. Following enzymatic digestion, samples were centrifuged at 400 × g for 1 min at 25 °C. The supernatant was removed, and the pellet was resuspended in 500 μl of W5 buffer containing 154 mM NaCl, 125 mM CaCl_2_, 5 mM KCl, 5 mM glucose and 2 mM MES (pH 5.7). After centrifugation at 400 × g for 1 min, the pellet was resuspended in 200 μl of MMG solution containing 0.4 M mannitol, 15 mM MgCl2 and 4 mM MES (pH 5.7).

For polyethylene glycol (PEG)-mediated transfection, 1–2 μl of plasmid DNA was mixed with 50 μl of the protoplast suspension and 50 μl of PEG solution containing 40% (w/v) PEG 6000, 100 mM CaCl_2_ and 0.2 M mannitol. After incubation for 5 min, 300 μl of W5 buffer was added, and the samples were centrifuged at 400 × g for 1 min at 25 °C. The pellet was resuspended in 200 μl of W5 buffer, and transfected protoplasts were incubated at 23 °C for 12–15 h before imaging with a confocal laser-scanning microscope (STELLARIS 8; Leica Microsystems). Fluorescence intensity profiles along the indicated ROIs were generated using Leica Application Suite X (LAS X; Leica Microsystems).

For the observation of PKS4-GFP, confocal microscopy images were taken with an Airy confocal microscope (Zeiss), model LSM 880. The samples *PKS4p::PKS4-GFP* and *PKS4p::mPKS4-D1-GFP* were excited with an Argon laser (488nm) and detection was done between 495 and 518 nm. In all cases a channel was set to detect chlorophyll and OLE1-RFP, exciting with the laser used for excitation of the fluorophore of interest, and detecting between 607 and 691 nm.

### Protein expression and purification

DNA fragments encoding RLD2-BRX (residues 1006–1066), AthBG1-Motif D (residues 320–340), PKS1-Motif D (residues 273–293), and BASL (residues 109–121) were amplified by PCR and cloned into pET47b(+) (Merck Millipore) or pGEX vectors (Cytiva). All plasmids were verified by Sanger sequencing and transformed into *Escherichia coli* Rosetta2 (DE3) cells (Merck Millipore).

For purification of RLD2-BRX, cells expressing N-terminally His-tagged RLD2-BRX were cultured in Luria– Bertani medium at 20 °C and induced with 0.1 mM isopropyl β-D-1-thiogalactopyranoside (IPTG) for 16–24 h. Cells were harvested, resuspended in buffer A [20 mM Tris-HCl (pH 8.0), 150 mM NaCl], and disrupted by sonication on ice. Lysates were clarified by centrifugation at 25,000 × g for 30 min at 4 °C. The supernatant was applied to complete His-Tag Purification Resin (Roche). After washing with buffer A containing 20 mM imidazole, bound proteins were eluted with buffer A containing 250 mM imidazole. The N-terminal His tag was removed by treatment with HRV 3C protease at 4 °C for approximately 10 h. RLD2-BRX was further purified by size-exclusion chromatography using a Superdex 75 pg column (Cytiva) equilibrated with buffer A.

For purification of AthBG1-Motif D, PKS1-Motif D and BASL peptides, cells expressing N-terminally GST-tagged peptides were cultured in Luria–Bertani medium at 37 °C and induced with 0.5 mM IPTG for 3–4 h. Cells were harvested, resuspended in buffer A, disrupted by sonication, and clarified by centrifugation. The supernatant was applied to glutathione Sepharose 4B resin (Cytiva). After washing with buffer A, GST-fusion proteins were eluted with buffer A containing 20 mM reduced glutathione. The GST tag was removed by treatment with HRV 3C protease at 4 °C for approximately 10 h. Peptides were further purified by size-exclusion chromatography using a Superdex 75 pg column (Cytiva), followed by reverse-phase chromatography on a Resource RPC column (Cytiva) using a linear gradient of 10–90% acetonitrile. Eluted peptide fractions were collected, dried using a vacuum concentrator, and stored at −30 °C until use. For crystallization, RLD2-BRX was co-expressed with AthBG1-Motif D (residues 320–340), PKS1-Motif D (residues 273–293), or BASL (residues 109–126) in *E. coli* Rosetta2 (DE3) cells. Cells were cultured in Luria– Bertani medium at 20 °C and induced with 0.1 mM IPTG for 16–24 h. Cell lysates were prepared as described above and applied to glutathione Sepharose 4B resin. Following washing with buffer A, bound proteins were eluted with buffer A containing 20 mM reduced glutathione and treated with HRV 3C protease at 4 °C for approximately 10 h. Protein complexes were further purified by size-exclusion chromatography using a Superdex 75 pg column (Cytiva) equilibrated with buffer A.

### Immunoblot analysis of PKS4–3×HA expression

Western blots to determine expression of the different *pPKS4:PKS4:3xHA* lines: Total proteins extracts, 80µl 2× Laemmli buffer for 20mg fresh weight seedlings, crushed with a pestle and immediately heated at 95°C for 5 min. Proteins were separated on 4-15% precast polyacrylamide gels (Bio-Rad) and transferred onto nitrocellulose using the Trans Blot Turbo RTA Transfer Kit. Anti-HA HRP monoclonal 3F10 was used at 1/4000 (12013819001, Roche), and anti-DET3 was used at 1/20000 dilutions (16) in 1X PBS containing 0.1% Tween-20 and 5% non-fat milk. Chemiluminescence signals were generated using Immobilon Western HRP Substrate (Millipore). Signals were detected with a Fujifilm ImageQuant LAS 4000 mini CCD camera system.

### Crystallization and data collection

Initial crystallization screening was performed using a Mosquito crystallization robot (TTP Labtech) and commercially available JCSG Core Suites I–IV and PACT Suite screens (QIAGEN).

Crystals of the RLD2-BRX–AthBG1-Motif D complex were obtained at 20 °C from protein complex at approximately 20 mg ml−1 and a reservoir solution containing 0.1 M HEPES–NaOH (pH 7.0), 0.2 M calcium acetate and 18% (w/v) polyethylene glycol (PEG) 8000. Crystals of the RLD2-BRX–PKS1-Motif D complex were obtained at 20 °C from protein complex at approximately 20 mg ml−1 and a reservoir solution containing 0.1 M Bis-Tris propane (pH 6.5), 0.2 M sodium formate and 20% (w/v) PEG 3350. Crystals of the RLD2-BRX– BASL complex were obtained at 20 °C from protein complex at approximately 24 mg ml−1 and a reservoir solution containing 0.1 M trisodium citrate (pH 5.0) and 8–10% (w/v) PEG 6000.

Crystals were transferred stepwise into cryoprotectant solutions containing 30% (v/v) ethylene glycol for the RLD2-BRX–AthBG1-Motif D and RLD2-BRX–BASL complexes, or 40% (w/v) PEG 3350 for the RLD2-BRX– PKS1-Motif D complex, and were flash-cooled in liquid nitrogen. X-ray diffraction data were collected at 100 K and a wavelength of 1.000 Å on beamlines BL41XU or BL45XU at SPring-8. Diffraction data were indexed, integrated and scaled using XDS. Crystallographic data collection statistics are provided in Extended Table 2.

### Structure determination and refinement

The structures of the RLD2-BRX complexes were determined by molecular replacement using the RLD2-BRX–LZY3-CCL complex structure (PDB ID: 6L0V) as the search model. Molecular replacement was performed using Phaser. Model building and refinement were carried out through iterative cycles of manual model building in Coot and refinement in PHENIX. Refinement statistics are summarized in Extended Table 2. Coordinates and structure factors for the RLD2-BRX–AthBG1-Motif D, RLD2-BRX–PKS1-Motif D and RLD2-BRX–BASL complexes have been deposited in the Protein Data Bank under accession codes 46PV, 46PW and 46PX, respectively.

### Structure and sequence comparisons

Pairwise structural comparisons were performed using the DALI server based on Cα-atom positions. Structure-based sequence alignments were generated from the superimposed structures. Molecular graphics were prepared using PyMOL Molecular Graphics System, version 1.7 (Schrödinger).

### Isothermal titration calorimetry

Isothermal titration calorimetry (ITC) was performed using an iTC200 calorimeter (Malvern Panalytical) at 20 °C. RLD2-BRX and peptide samples were prepared in buffer B containing 20 mM HEPES–NaOH (pH 7.0) and 150 mM NaCl. For BASL measurements, buffer B was supplemented with 10% (v/v) dimethyl sulfoxide (DMSO).

For the AthBG1-Motif D measurement, 2.5 μl aliquots of 500 μM AthBG1-Motif D peptide (residues 320–340) were injected into 50 μM RLD2-BRX. For the PKS1-Motif D measurement, 2.5 μl aliquots of 554 μM PKS1-Motif D peptide (residues 273–293) were injected into 50 μM RLD2-BRX. For the BASL measurement, 2.5 μl aliquots of 291 μM BASL peptide (residues 109–121) were injected into 30 μM RLD2-BRX.

Heat changes arising from peptide dilution were subtracted from the binding isotherms. Data were fitted using a one-site binding model with ORIGIN software supplied with the instrument.

### Pull-down binding assays

All point mutations were introduced by site-directed mutagenesis. For in vitro pull-down assays, purified GST-fusion peptides were immobilized on glutathione Sepharose 4B resin in buffer A [20 mM Tris-HCl (pH 8.0), 150 mM NaCl] and incubated with purified RLD2-BRX at 4 °C. After extensive washing with buffer A, bound proteins were eluted and analyzed by SDS–PAGE.

The relative amount of RLD2-BRX recovered in the bound fraction was quantified from three independent experiments. Error bars represent the standard deviation.

### Phylogenetic and comparative analyses of RLD, GNOM and RLD-interacting protein families

Protein sequences of RLD, GNOM, LZY, PKS, AthBG and BASL family members from representative green plant species were identified by BLASTP searches against the National Center for Biotechnology Information (NCBI) protein database. Arabidopsis thaliana RLD3, GNOM, LZY1, LZY3, PKS1, AthBG1 and BASL protein sequences were used as queries. Candidate homologues were assessed based on sequence similarity and conservation of characteristic functional domains.

Phylogenetic trees of RLD and GNOM proteins were constructed from amino acid sequence alignments using MEGA11 (33). Evolutionary history was inferred using the neighbour-joining method. Node support was assessed by bootstrap analysis with 2,000 replicates. Evolutionary distances were calculated using the Poisson correction method and are expressed as the number of amino acid substitutions per site. All positions containing gaps or missing data were excluded by complete deletion. Trees were drawn to scale, with branch lengths corresponding to evolutionary distances. For comparative presence profiling, homologues with high sequence similarity were scored as present, whereas proteins with low overall sequence similarity but conservation of characteristic functional domains were classified separately.

---

**Extended Fig. 1.**
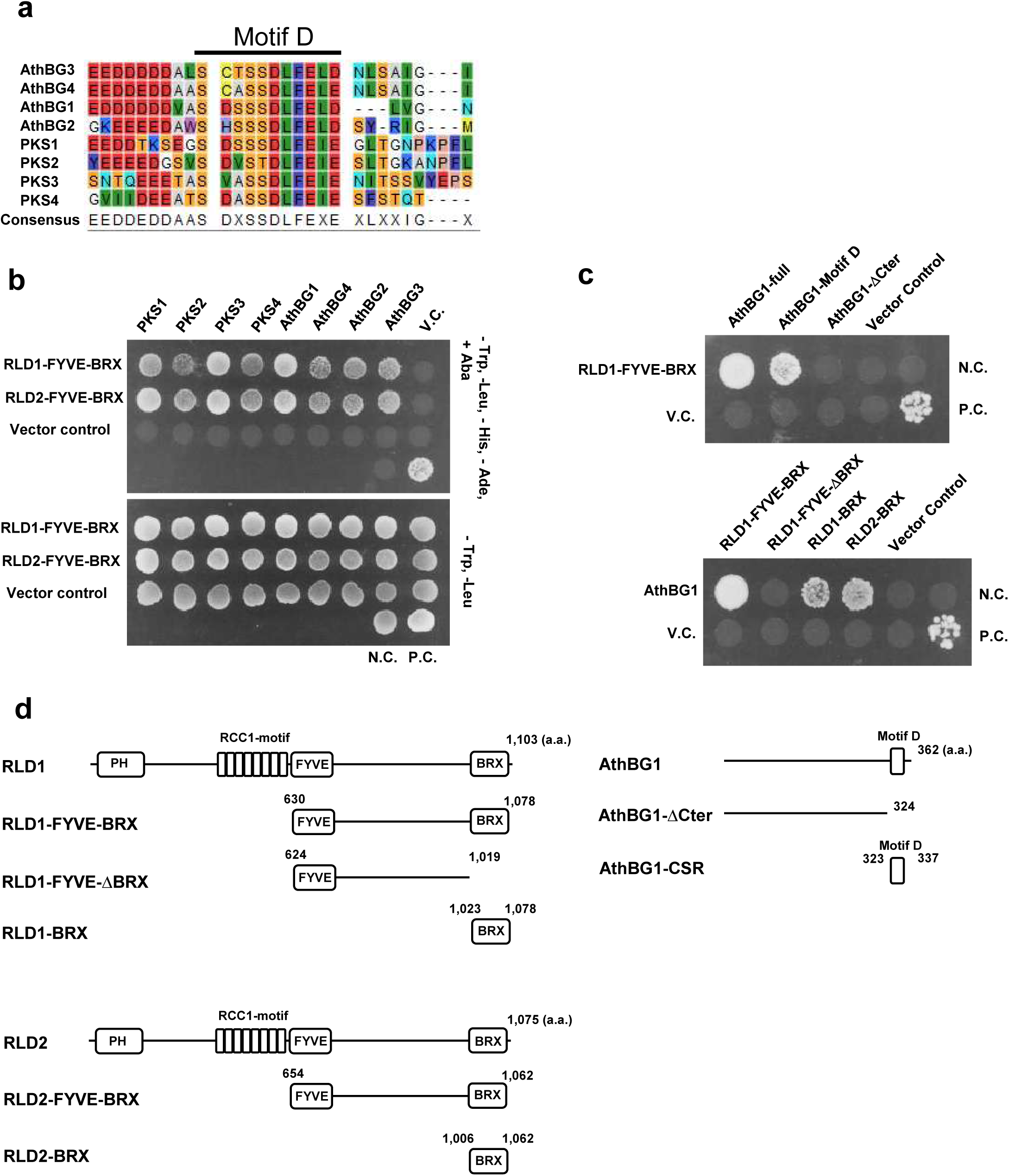
Interaction of RLD-FYVE-BRX with PKS and BG family proteins. **(a)** Highly conserved motif D sequences in *Arabidopsis* BG and PKS family proteins. **(b)** RLD1-FYVE-BRX and RLD2-FYVE-BRX interact with PKS and BG family proteins in a yeast two-hybrid (Y2H) assay. Prey constructs expressing RLD1-FYVE-BRX (830–1078 aa) and RLD2-FYVE-BRX (864–1082 aa) were fused to the activation domain (AD), while bait constructs expressing full-length PKS and BG family proteins were fused to the DNA-binding domain (BD). The interaction between T-antigen and p53 served as a positive control (P.C.), and that between T-antigen and human lamin as a negative control (N.C.). Diploid yeast cells were grown on synthetic medium lacking Trp, Leu, His, and Ade (–Trp/–Leu/–His/–Ade) to test for interaction. **(c)** Interaction between RLD1-FYVE-BRX and truncated AthBG1, and between AthBG1 and truncated RLD1, analyzed using the Y2H system. **(d)** Schematic representation of protein fragments used in the Y2H assays.

**Extended Fig. 2.**
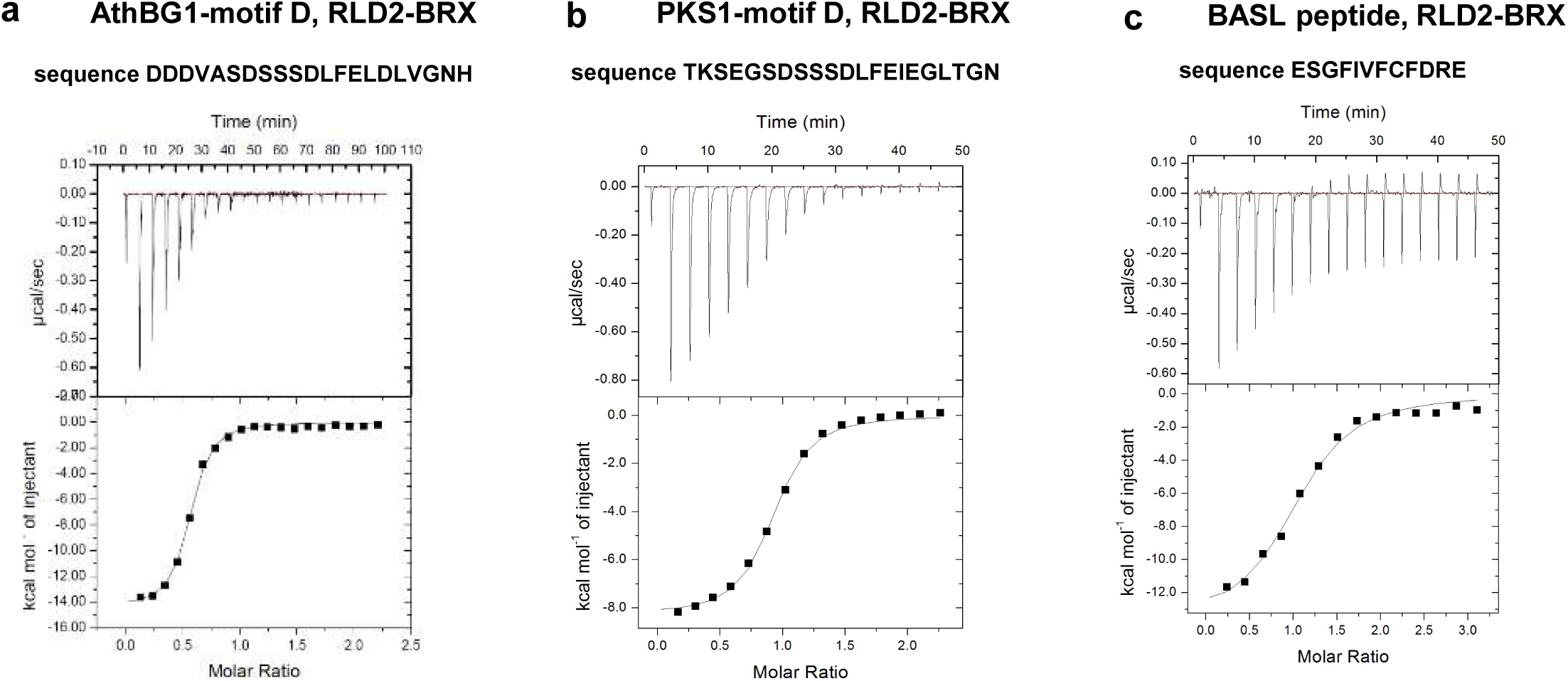
ITC analysis of BRX interactions.(a) Binding of the RLD2-BRX domain to the AthBG1 motif-D peptide. The ITC profile indicates formation of a 1:1 complex with *K*ᴅ = 820 ± 95 nM. Δ*H*, Δ*S*, and *N* are –14.23 ± 0.21 kcal mol⁻¹, –21.1 cal mol⁻¹ deg⁻¹, and 0.53 ± 0.006, respectively. **(b)** Binding of the RLD2-BRX domain to the PKS1 motif-D peptide (*K*ᴅ = 1.23 ± 0.14 μM). Δ*H*, Δ*S*, and *N* are – 8.28 ± 0.16 kcal mol⁻¹, –1.21 cal mol⁻¹ deg⁻¹, and 0.88 ± 0.009, respectively. **(c)** Binding of the RLD2-BRX domain to the BASL peptide (*K*ᴅ = 1.83 ± 0.35 μM). Δ*H*, Δ*S*, and *N* are –1.38 ± 0.63 kcal mol⁻¹, –20.6 cal mol⁻¹ deg⁻¹, and 1.04 ± 0.035, respectively.

**Extended Fig. 3.**
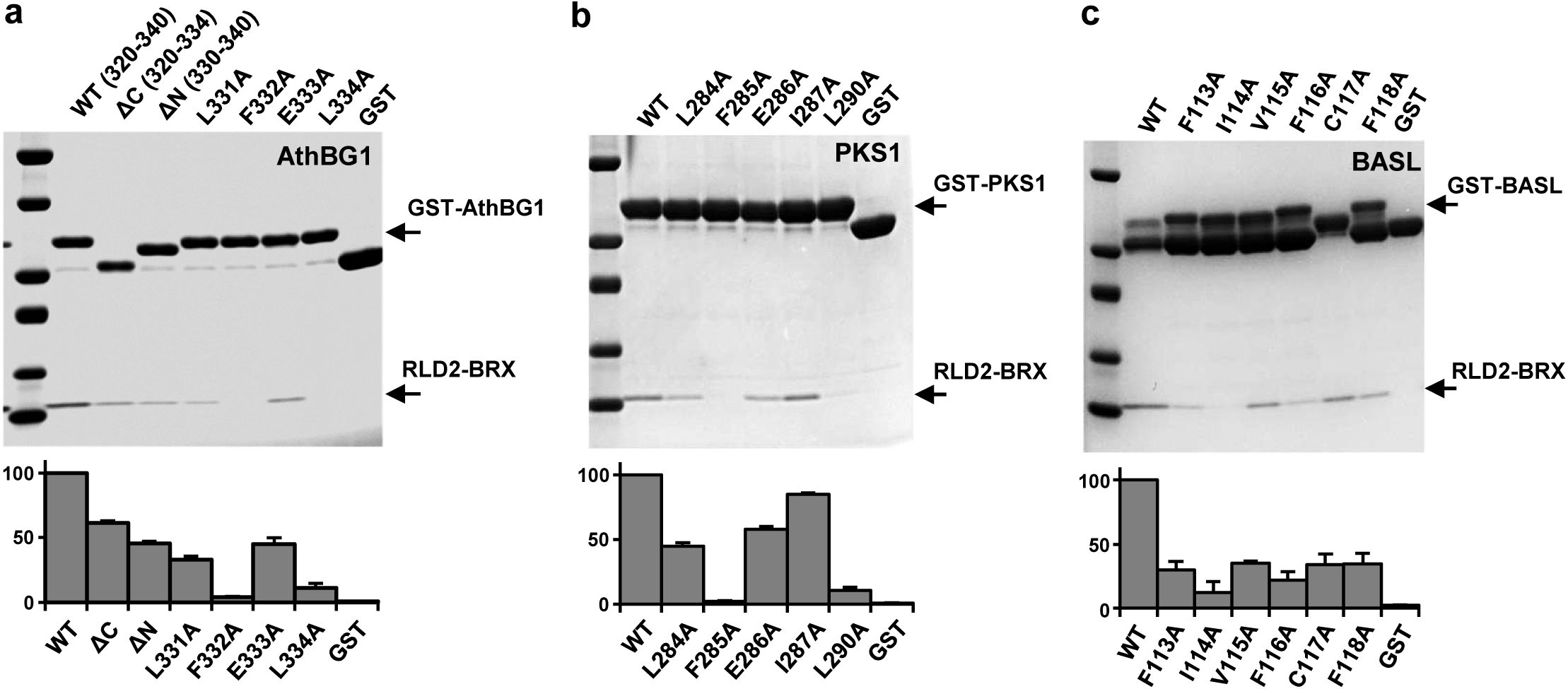
Functional validation of BRX–partner interactions. (a–c) Pull-down assays showing binding between GST–RLD2-BRX and GST-tagged peptides: (a) PKS1-motif D, (b) AthBG1-motif D, and (c) BASL peptide. Top panels, SDS–PAGE analysis; bottom panels, quantification of pulled-down RLD2-BRX.

**Extended Fig. 4.**
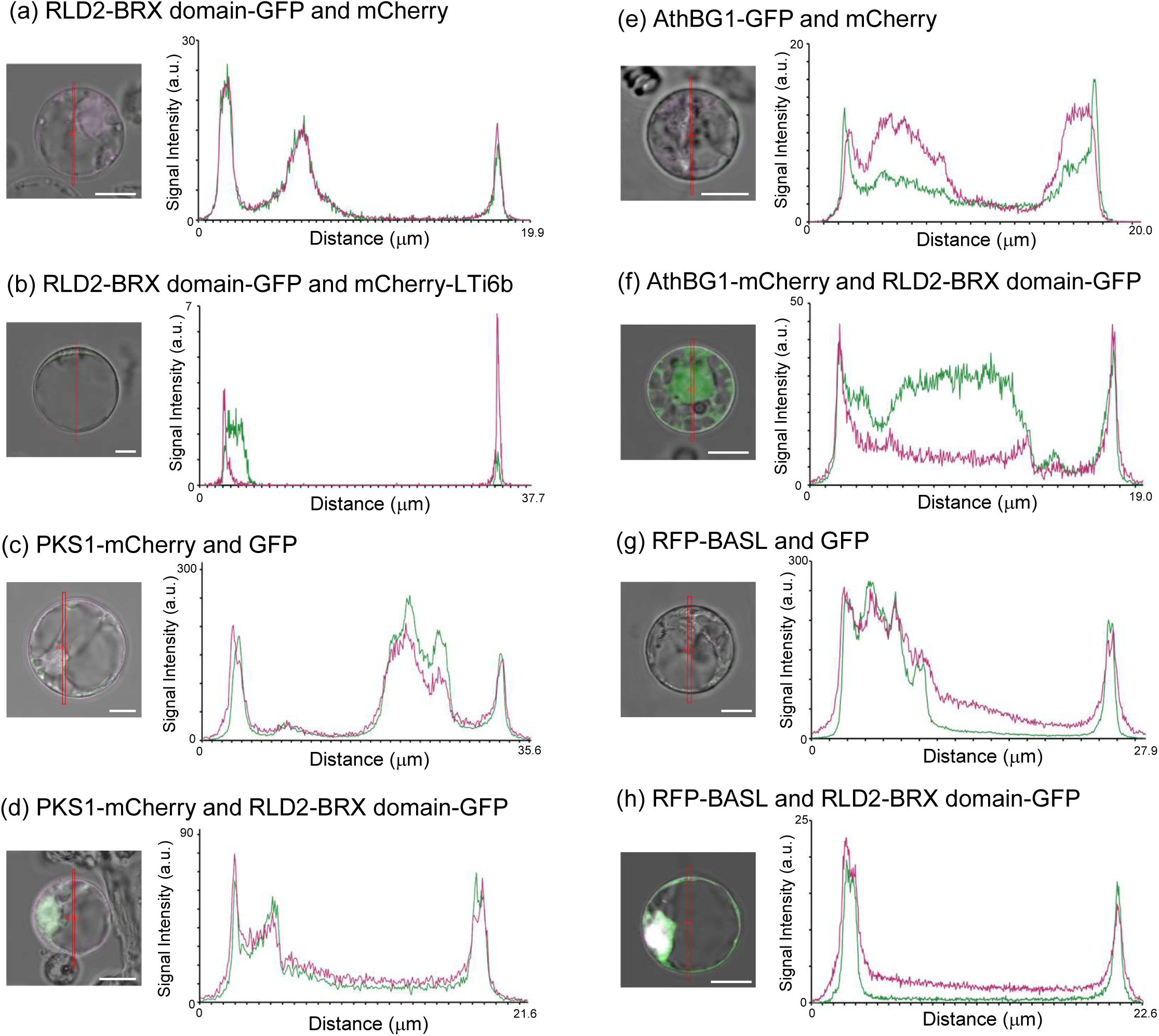
Fluorescence intensity profiles of co-localization assays. Fluorescence intensity plots along ROIs from the transient expression assay shown in Fig. 3. Green lines indicate GFP signals; magenta lines indicate mCherry or mRFP signals. Scale bar = 7.5 µm

**Extended Fig. 5.**
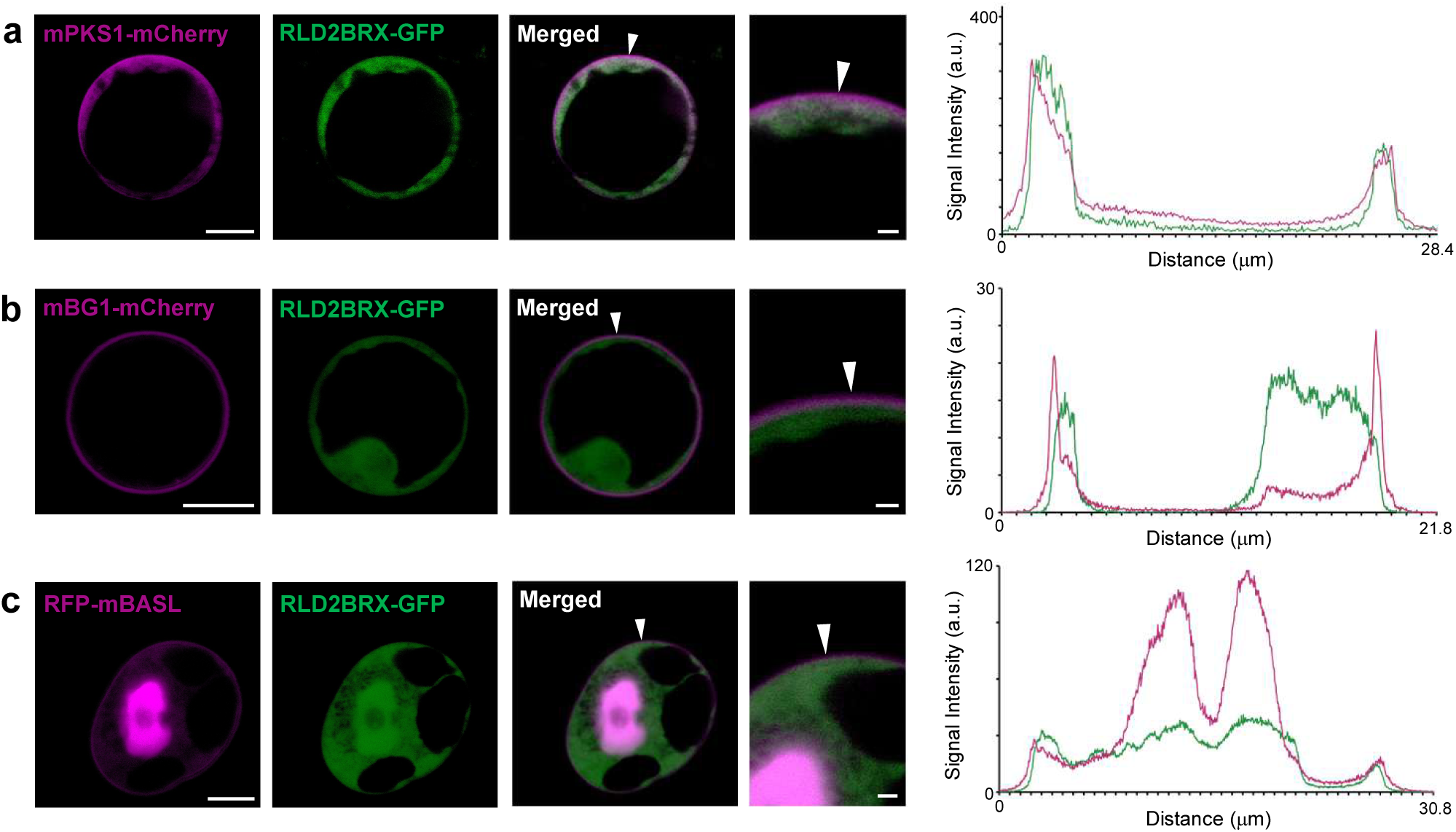
Mutational validation of BRX-dependent RLD recruitment by PKS1, AthBG1, and BASL. **(a–c)** Co-localization analysis of mutated PKS1, AthBG1, and BASL with the RLD2-BRX domain in transiently transformed *Arabidopsis* root protoplasts. Guided by crystal structure and pull-down results, mutations were introduced into residues critical for BRX interaction. Co-expression of transgenes: (a) mutated PKS1–mCherry with RLD2-BRX domain–GFP; (b) mutated AthBG1–mCherry with RLD2-BRX domain–GFP; (c) mutated BASL–RFP with RLD2-BRX domain–GFP. Arrowheads indicate regions used for intensity plots. Scale bar = 7.5 µm

**Extended Fig. 6.**
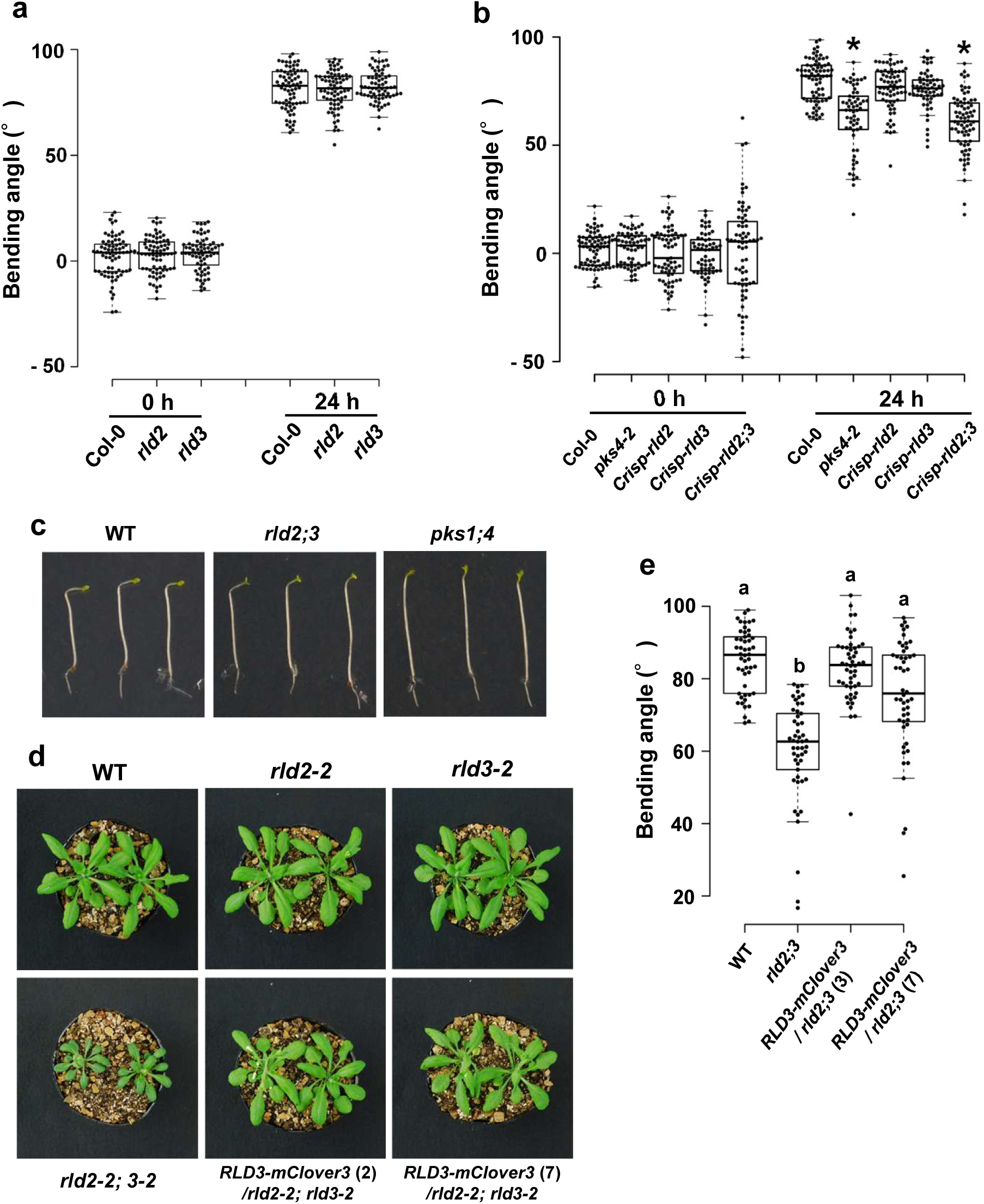
Phototropic responses of *rld* mutants and complementation analysis. **(a)** Phototropism of etiolated hypocotyls in *rld2* and *rld3* mutants. Seedlings were irradiated with unilateral blue light (0.1 μmol m⁻² s⁻¹), and hypocotyl bending was measured after 24 h (*n* = 70–75). Box plots show medians, 25th–75th percentiles, whiskers at 1.5× IQR, and outliers as dots. No significant differences were detected compared with WT (one-way ANOVA, Dunnett’s test). **(b)** Phototropism of *rld2*, *rld3*, and *rld2 rld3* double mutants generated by CRISPR–Cas9. Conditions as in (a). *n* = 58–74. Asterisks indicate significant differences from WT (*P* < 0.05, Dunnett’s test). **(c)** Representative images of phototropic responses in *rld2 rld3* and *pks1 pks4* double mutants, taken 12 h after the onset of unilateral blue-light irradiation. **(d)** Dwarf phenotype of *rld2*, *rld3*, and *rld2 rld3* double mutants in rosette leaves, and phenotypic complementation by *RLD3–mClover3*. **(e)** Complementation of the *rld2 rld3* double mutant phototropic defect by *RLD3–mClover3*. Hypocotyl curvature was measured 24 h after unilateral blue light (0.1 μmol m⁻² s⁻¹). *n* = 50. Different letters indicate significant differences (Tukey–Kramer test, *P* < 0.01).

**Extended Fig. 7.**
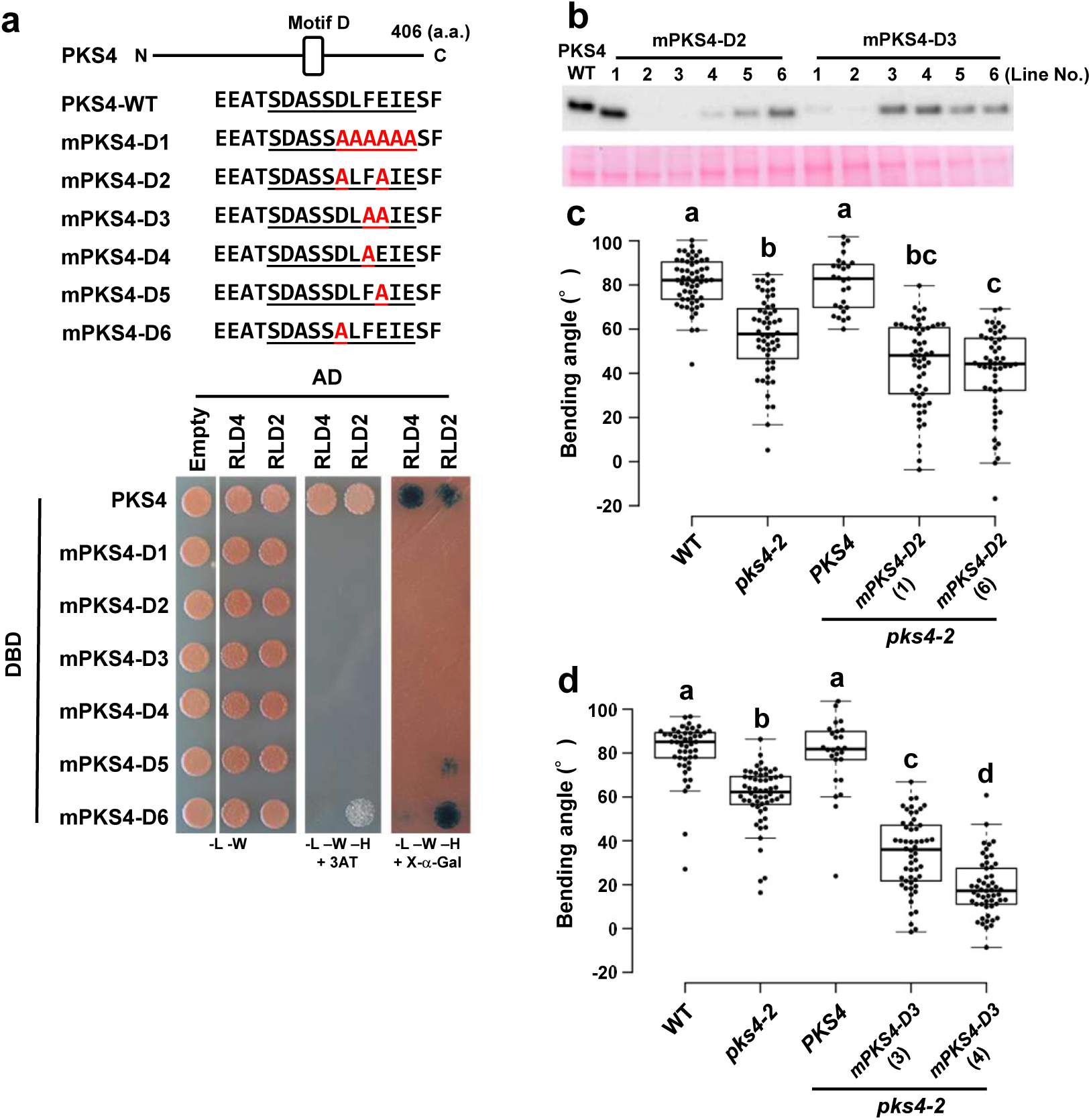
Functional analysis of PKS4 motif D in BRX interaction and phototropism. **(a)** Effect of various motif D mutations in PKS4 on interaction with RLD2 and RLD4 in the Y2H assay. Variants of PKS4 motif D are illustrated schematically (top). Prey constructs expressing RLD2 and RLD4 were fused to AD, and bait constructs expressing WT or mutant PKS4 (mPKS4-D1–D5) were fused to BD. Diploid yeast cells were grown on –Trp/–Leu/–His + 3-AT and –Trp/–Leu/–His + X-α-Gal media to test interaction. **(b–d)** Effect of motif D mutations on hypocotyl phototropism. GFP-tagged WT, mPKS4-D2 and mPKS4-D3 were expressed in the *pks4-2* mutant. Lines with confirmed protein expression (Western blot) were analyzed for phototropic bending under unilateral blue light (0.1 μmol m⁻² s⁻¹). Hypocotyl curvature was measured after 24 h (*n* = 27–57 for mPKS4-D2; *n* = 26–56 for mPKS4-D3). Different letters indicate significant differences (Tukey–Kramer test, *P* < 0.01).

**Extended Fig. 8.**
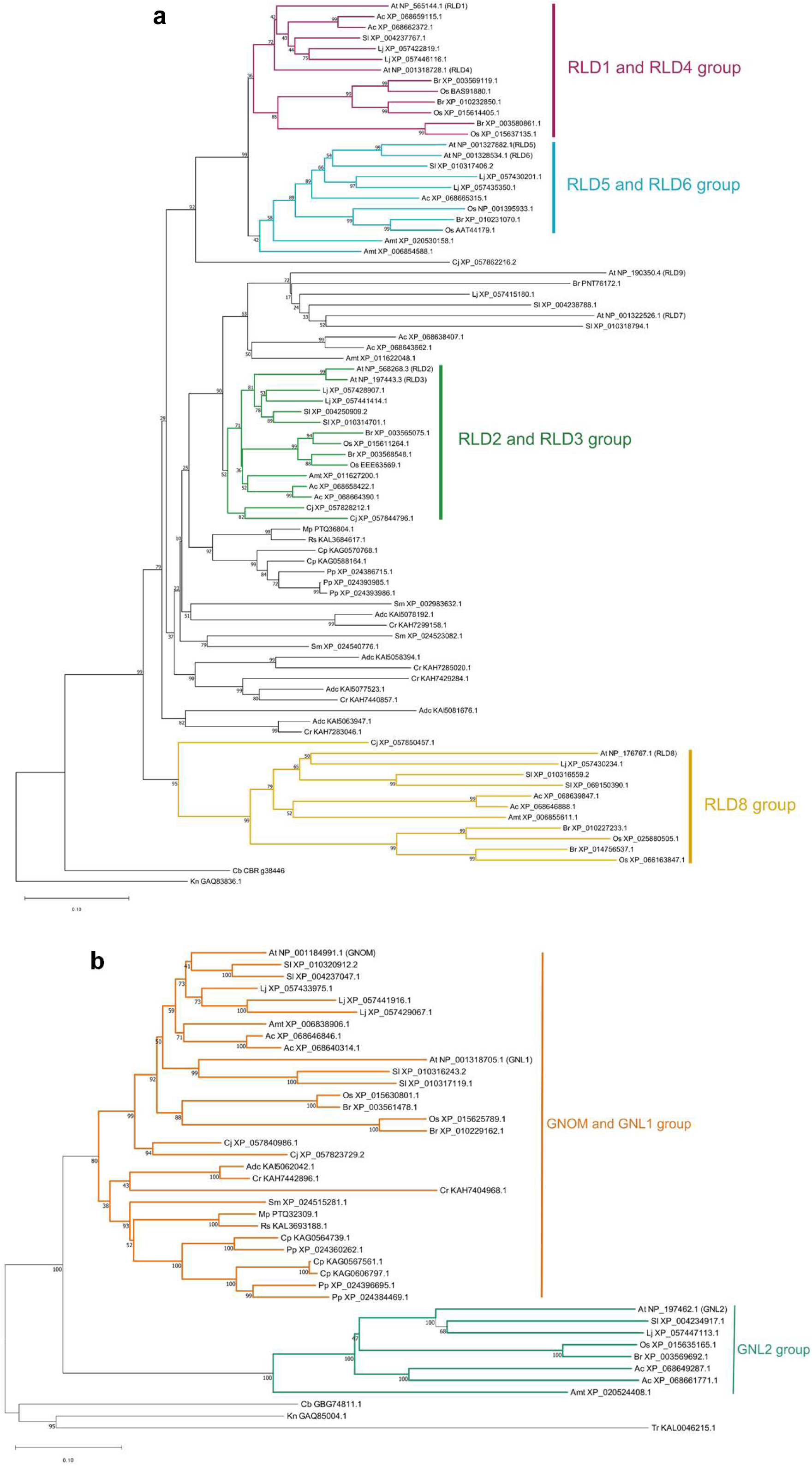
Phylogenetic analysis of RLD and GNOM families. Phylogenetic trees of. **(a)** RLD and (b) GNOM proteins inferred using the neighbor-joining method. Bootstrap support values (2000 replicates) are shown next to branches. Scale bars indicate the number of amino acid substitutions per site. Evolutionary distances were computed using the Poisson correction method, and analyses were conducted in MEGA11. Abbreviations: Ac, *Aristolochia californica*; Adc, *Adiantum capillus-veneris*; Amt, *Amborella trichopoda*; At, *Arabidopsis thaliana*; Br, *Brachypodium distachyon*; Cb, *Chara braunii*; Cj, *Cryptomeria japonica*; Cp, *Ceratodon purpureus*; Cr, *Ceratopteris richardii*; Kn, *Klebsormidium nitens*; Lj, *Lotus japonicus*; Mp, *Marchantia polymorpha*; Os, *Oryza sativa*; Pp, *Physcomitrium patens*; Rs, *Riccia sorocarpa*; Sl, *Solanum lycopersicum*; Sm, *Selaginella moellendorffii*; Tr, *Trebouxia* sp.

**Extended Table. 1.** Interacting proteins identified by yeast-two-hybrid screening using RLD1-FYVE-BRX as bait.

| Counts | Identified proteins | AGI code | Motif D | CCL |
| --- | --- | --- | --- | --- |
| 23 | LZY1 | At5G14090 | No | Yes |
| 17 | PKS2 | At1G14280 | Yes | No |
| 15 | xylem cysteine proteinase 2 | At1G20850 | No | No |
| 14 | RING-H2 finger protein ATL72 | At3G10910 | No | No |
| 10 | AthBG3 | At1G54200 | Yes | No |
| 9 | transcription factor MYB3 | At1G22640 | No | No |
| 6 | putative 30S ribosomal subunit S11 protein | At1G31817 | No | No |
| 6 | uncharacterized protein | At3G50340 | No | No |
| 5 | AthBG1 | At5G12050 | Yes | No |
| 5 | chaperone DnaJ-domain superfamily protein | At1G79030 | No | No |
| 5 | uncharacterized protein | At5G67020 | No | No |
| 5 | myb domain protein 7 | At2G16720 | No | No |
| 3 | AthBG2 | At3G42800 | Yes | No |
| 2 | LZY6 | At3G27025 | No | Yes |
| 1 | LZY3 | At1G72490 | No | Yes |
| 1 | PKS3 | At1G18810 | Yes | No |

**Extended Table. 2.** X-ray data collection and refinement statistics.

|  | RLD2-AthBG1 | RLD2-PKS2 | RLD2-BASL |
| --- | --- | --- | --- |
| Data collection |  |  |  |
| Space group | *P*321 | *P*2<sub>1</sub>2<sub>1</sub>2<sub>1</sub> | *P*2<sub>1</sub>2<sub>1</sub>2<sub>1</sub> |
| Cell dimention |  |  |  |
| *a*, *b*, *c* (Å) | 82.4, 82.4, 33.3 | 20.6, 51.7, 132.5 | 45.2, 50.5, 71.9 |
| $\alpha$, $\beta$, $\gamma$ (°) | 90.0, 90.0, 120 | 90.0, 90.0, 90.0 | 90.0, 90.0, 90.0 |
| Resolution (Å)* | 41.22-2.00 (2.03-2.00) | 48.13-1.80 (1.84-1.80) | 45.23-1.52 (1.55-1.52) |
| R<sub>meas</sub> (%)* | 4.0 (34.8) | 9.6 (69.3) | 4.9 (82.0) |
| CC (1/2) | 1.00 (0.99) | 1.00 (0.94) | 1.00 (0.85) |
| *I* / $\sigma$ (*I*)* | 30.9 (7.3) | 11.7 (2.4) | 18.9 (2.0) |
| Completeness (%)* | 98.2 (98.2) | 100.0 (100.0) | 99.9 (99.9) |
| Redundabcy* | 9.8 (10.0) | 10.6 (9.7) | 5.5 (4.6) |
| Mosaicity | 0.20 | 0.08 | 0.35 |
| Refinement |  |  |  |
| Resolution (Å) | 41.22-2.00 | 48.13-1.80 | 41.34-1.52 |
| No. reflections | 8790 | 13871 | 25946 |
| R<sub>work</sub> / R<sub>free</sub> (%) | 21.7 23.4 | 19.3/ 21.4 | 19.1/ 19.7 |
| No. atoms |  |  |  |
| Protein | 585 | 1201 | 1149 |
| Water | 48.97 | 29.37 | 37.70 |
| Ligand | 59.05 |  | 46.05 |
| R.m.s. deviations |  |  |  |
| Bond lengths (Å) | 0.005 | 0.009 | 0.06 |
| Bond angles (° ) | 0.723 | 0.956 | 0.824 |

One crystal was used for each data set. *R*<sub>free</sub> was calculate on a random 5% reflections of tha data.
\*Highest resolution shell is shown in parenthesis.

**Supplementary Table. 1.**
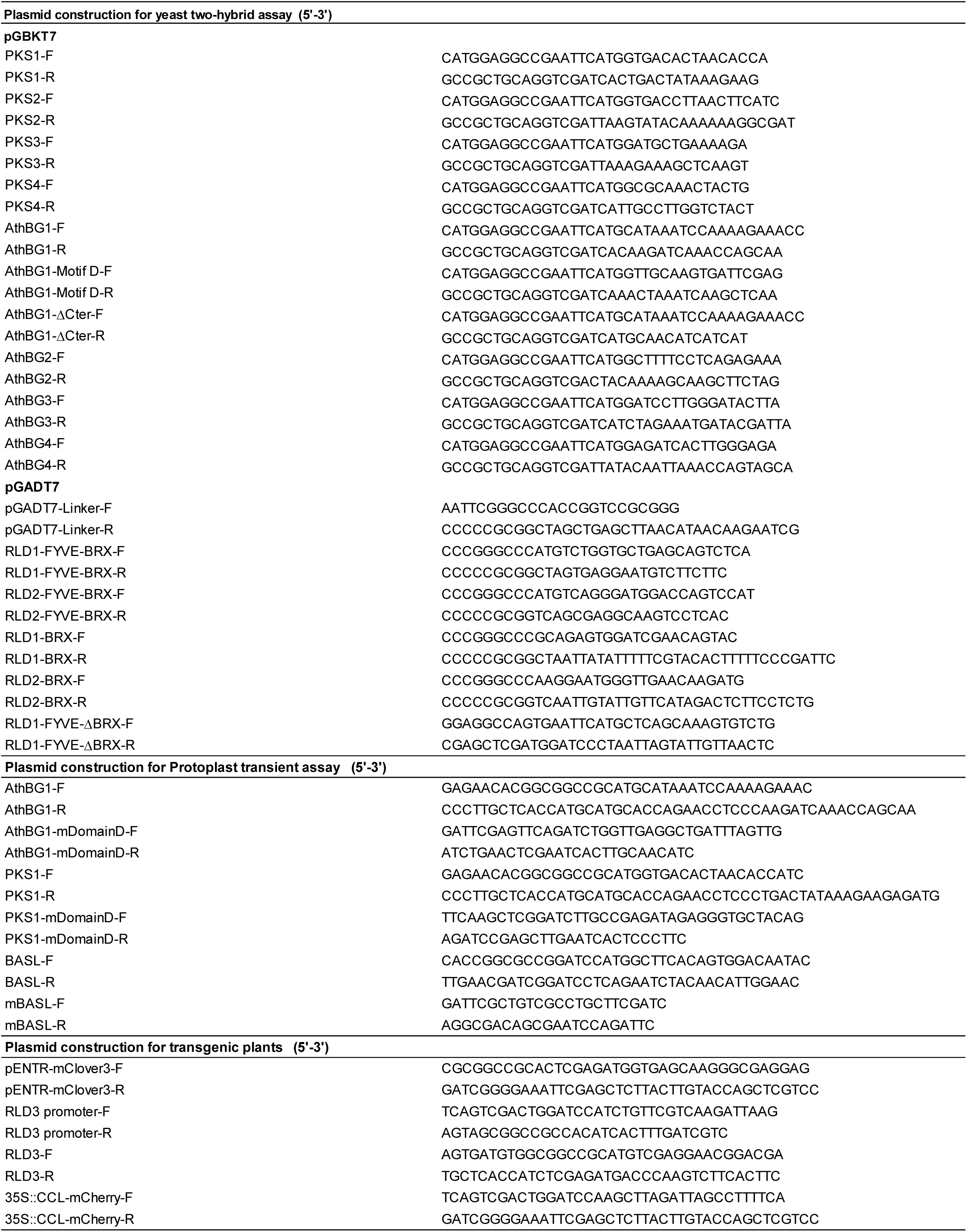
Primer used in this study.

